# Mllt11 regulates the migration and neurite outgrowth of cortical projection neurons during development

**DOI:** 10.1101/2021.10.31.466682

**Authors:** Danielle Stanton-Turcotte, Karolynn Hsu, Samantha A. Moore, Makiko Yamada, James P. Fawcett, Angelo Iulianella

**Author notes:** Molecular Pharmacology Program, Memorial Sloan Kettering Cancer Center, 1275 York Avenue, ZRC-1863, New York, NY 10021. **Contributions:** DST, KS, SM did the experiments, analysis and figures. DST and AI co-wrote the manuscript. JF advised with the GST-pull down experiments and assisted with the proteomic assays. AI assisted with imaging and supervised the project.

## Abstract

The formation of connections within the mammalian neocortex is highly regulated by both extracellular guidance mechanisms and intrinsic gene expression programs. There are generally two types of cortical projection neurons: those that project locally and interhemispherically, and those that project to sub-cerebral structures such as the thalamus, hindbrain, and spinal cord. The regulation of cortical projection morphologies is not yet fully understood at the molecular level. Here we report a role for Mllt11 (Myeloid/lymphoid or mixed-lineage leukemia; translocated to chromosome 11/All1 Fused Gene From Chromosome 1q) in the migration and neurite outgrowth of callosal projection neurons during brain formation. We show that Mllt11 expression is exclusive to developing neurons and is enriched in the developing cortical plate, particularly during the formation of the upper or superficial cortical layers. In cultured primary cortical neurons, Mllt11 is detected in varicosities and growth cones as well as the soma. Using conditional loss-of-function and gain-of-function analysis we show that Mllt11 is a required for neuritogenesis and proper migration of upper layer cortical projection neurons. Loss of *Mllt11* in the superficial cortex leads to a severe reduction in fibres crossing the corpus callosum, a progressive loss in the maintenance of upper layer projection neuron gene expression, and dysplasia of dendritic arborisation patterns. Proteomic analysis revealed that Mllt11 associates with cytoskeletal components including stabilized microtubules consistent with a role in neuronal migration and neuritogenesis. Taken together, our findings support a role for Mllt11 in promoting the formation of mature projection neuron morphologies and connectivity in the cerebral cortex.

## INTRODUCTION

The mammalian neocortex is a complex structure, underlying the capacity for executive function, sensory processing, emotion, motor output, and cognition. It is a laminated structure composed of six molecularly and morphologically distinct layers populated by a heterogeneous collection of excitatory pyramidal cortical projection neurons (CPNs) and inhibitory interneurons (INs). This anatomical organization emerges from the tight regulation of neurogenesis coupled with migration of newborn neurons to their final laminar position, where they acquire distinct molecular identities and morphological traits necessary to form functional circuitry within the brain. A fundamental question concerns how the migration of newborn neurons is controlled in the sequential generation of more superficial cortical layers and their subsequent connectivity patterns. While it is clear that the cytoskeleton plays a key role in neuronal translocation, how it impacts CPN subtype specific neurite morphogenesis and the maintenance of laminar transcriptional programs it not well understood.

The sorting of differentiating CPN subtypes into discrete, functional layers occurs as cells migrate radially from progenitor domains located along the ventricular zone (VZ) and adjacent subventricular zone (SVZ) along the apicobasal axis to invade the cortical plate (CP). Early in cortical development newly-born neurons migrate along radial glial fibres projecting from the basal surface of VZ progenitors to the pia via somal translocation (Miyata et al., 2001). Forces exerted on the cytoskeleton, including the microtubules that form the perinuclear cage, couple with motor proteins to enable appropriate mechanotransduction to push and pull the nucleus toward the pia until a neuron reaches its terminal location (Bellion et al., 2005; Tsai et al., 2007). As the somas of these cells reach the VZ, previously anchored apical junctional complexes are downregulated, releasing the soma from the VZ, and the trailing apical pole of the cells becomes the nascent axon projecting to form white matter tracts (Ayala et al., 2007; Marin and Rubenstein, 2003; Nadarajah et al., 2001; Sakakibara et al., 2014). Consequently, the intermediate zone (IZ) and CP become more densely populated with neurons and projections, and multiple modes of motility must be employed by later-born neurons to overcome the previously deposited neuronal layers. These events also rely on coordinated cytoskeletal reorganization as cells transiently acquire multipolarity to weave through the dense IZ and then reacquire bipolarity upon invasion of the CP via somal translocation (Sakakibara et al., 2014). The growth of axons during brain formation also involves dynamic cytoskeletal reorganization at the growth cone in response to external guidance cues or cell-cell contacts (Ayala et al., 2007; Hirokawa and Takemura, 2004).

Microtubule growth, retraction and stabilization are controlled by complex signaling from extracellular cues, and the coordination of these events are necessary for appropriate neuronal migration and neurite extension. The ability to couple developmental cues with construction or degradation of microtubule networks underlie the propensity of CPNs to form connections both intracortically and with other brain regions. As such, mutations of tubulin subtypes have been linked to neurodevelopmental disorders that affect cortical formation, stabilization, and projections. For instance, mutation of α-tubulin isoforms cause lissencephaly and other gyrification defects, as well as reduction in white matter resulting in abnormalities of the corpus callosum (CC) and internal capsule (IC) (Bahi-Buisson et al., 2008; Jansen et al., 2011; Keays et al., 2007; Kumar et al., 2010; Poirier et al., 2007; Tian et al., 2010). β-tubulin mutations have been linked with cortical dysplasia, epilepsy, decreased neuronal density, enlarged ventricles, white matter thinning, and defects in migration and radial glial fibre extension due to their role in axonal growth, guidance, cell-cycle-dependent proliferation (Breuss et al., 2012; Breuss et al., 2017; Cushion et al., 2014; Ejaz et al., 2017; Hersheson et al., 2013; Jaglin et al., 2009; Jamuar et al., 2014; Ngo et al., 2014; Poirier et al., 2010; Rodan et al., 2017; Tischfield et al., 2010; Whitman et al., 2016). Mutation of γ-tubulin has also been shown to cause a wide range of phenotypes common to other tubulin isoforms due to its role in stabilizing growth and nucleation of microtubules (Brock et al., 2018; Draberova et al., 2017; Ivanova et al., 2019; Poirier et al., 2013; Yuba-Kubo et al., 2005).

Much like tubulin, mutations of microtubule associated proteins (MAPs) including stabilizing or nucleating proteins, motor proteins, and cargo-specific adapter proteins that govern trafficking of organelles and proteins to specific regions of a neuron, have also been linked with various forms of neurodevelopmental disorders (Moffat et al., 2015; Moon and Wynshaw-Boris, 2013). Mutations of these proteins can also affect neuronal development at the level of mitosis, axon guidance, disturbance of fasciculation of axonal tracts, or impaired migration. Mutations and aberrant localization of neurofilaments mediated by MT-based transport have also been linked with schizophrenia, bipolar disorder (BPD), and autism spectrum disorders (ASD) (Clark et al., 2007; English et al., 2009; Kristiansen et al., 2006; Pennington et al., 2008; Sivagnanasundaram et al., 2007; Yao et al., 2021). A common thread between all mutations of tubulin isotypes and associated proteins is their propensity to give rise to phenotypes commonly associated with neurodevelopmental disorders through migration, cell fate specification, and axonal regulatory mechanisms required for functional connectivity within the cortex and between the cortex and other regions.

Although axonal regulatory mechanisms are conserved throughout the cortex, the targeting and wiring of each cortical layer is also dependent upon its unique molecular characteristics. For instance, each cortical layer expresses a unique program of transcription factors that provide the necessary molecular context crucial for synapses formation with the appropriate target regions of the brain. With the exception of the Cajal-Rietzus cells of layer 1 (L1), which are born first cells during corticogenesis and secrete Reelin to guide the radial migration of nascent neurons (Chai et al., 2009; Chai et al., 2016; Frotscher, 1998; Gil-Sanz et al., 2013), this process begins with Tbr1+ neurons which populate layer 6 (DL6) and project to the thalamus (Hevner et al., 2001; Thomson, 2010). Subsequently, the Bcl11b/Ctip2+ neurons of layer 5 (DL5) are born, which mostly connect to subcerebral regions (Arlotta et al., 2005). DL4 neurons, which receive input from the thalamus, can be identified by their co-expression of Foxp2 and Satb2 (Hisaoka et al., 2010; Leone et al., 2015; Lopez-Bendito and Molnar, 2003). The last cortical cell types generated are those of layers 2/3 (L2/3) in the superficial cortex, of which 80% project callosally and fewer than 20% project ipsilaterally within the cortex, identified by the expression of Cux1, Cux2 (Nieto et al., 2004; Zimmer et al., 2004), and Satb2 (Alcamo et al., 2008; Britanova et al., 2008; Leone et al., 2015).These transcription factors act in cross-repressive networks to regulate the organized chronological generation of cortical layers and functional diversity of projection neuron subtypes (Toma et al., 2014).

In an effort to identify regulators of neural migration and differentiation, we previously reported the expression of Mllt11/Af1q (Myeloid/lymphoid or mixed-lineage leukemia; translocated to chromosome 11/All1 Fused Gene From Chromosome 1q) in the developing central nervous system, including the cortical plate (Yamada et al., 2014). Mllt11 is a vertebrate-specific 90 amino acid protein that possesses a nuclear export signal but otherwise has poorly defined functional domains. Mllt11 was first identified in an infant with acute myelomonocytic leukemia carrying the t(1;11)(q21;q23) translocation that creates an abnormal protein fused to Mll (Tse et al., 1995). Mllt11 may play a normal role in neural differentiation, as its transcription is down-regulated by RE1 silencing transcription factor, a key factor involved in regulating terminal differentiation of neurons (Hu et al., 2015). However, functional studies addressing its role in neural development are currently lacking. Here we report that Mllt11 is a neural-specific tubulin-associating protein and its conditional inactivation in the superficial layers of the mouse brain results in improper migration and formation of callosal projections. Additionally we show that *Mllt11* loss led to dysplasia of the superficial cortex characterized by a severe loss of upper layer CPN dendritic morphologies resulting from neurite outgrowth defects. Interestingly, the inability to initiate and maintain functional projections in *Mllt11* mutant brains was accompanied by a loss of upper cortical layer (UL)-specific transcriptional programs. On the other hand, *Mllt11* overexpression promoted invasion of the cortical plate by differentiating neurons. Mechanistically, we provide evidence that Mllt11 interacts with stabilized microtubules, explaining the phenocopy of *Mllt11* mutants with brain tubulinopathies (Bahi-Buisson et al., 2014). Altogether, these findings contribute significantly to our understanding of the genetic regulation of projection neuron development, dendritic complexity, and axonal connectivity.

## RESULTS

### Conditional Knockout of Mllt11 from Upper Layer 2/3

Mllt11 is expressed in developing neurons of the central nervous system, including the neocortex, but its role in cortical neurogenesis is unknown (Yamada et al, 2014). To investigate this, we generated mice carrying a null mutation in the *Mllt11* gene using embryonic stem (ES) cell clones obtained from the mouse knockout consortium project (UCDavis KOMP repository, *Mllt11^tm1a(KOMP)Mbp^*). We used the targeted (unflipped) allele to visualize *Mllt11* by β-galactosidase (β-gal) staining due to the *LacZ* gene inserted into the *Mllt11* locus (Fig. S1A, S4A). Initial *Mllt11* locus activity begins in the pallial MZ at E12.5, showing stronger expression in the intermediate zone (IZ) of the fetal cortex at E14.5 coinciding with the birth of basal progenitors fated to become UL2/3 CPNs, intensifying in the superficial cortical layers by E16.5-E18.5 (Fig S4A). This confirms our previous report of *Mllt11* mRNA and protein localization in these regions (Yamada et al., 2014). Subsequent in situ hybridization revealed that cortical expression is maintained until postnatal day 21 (P21) when it begins to taper to levels indistinguishable from background at P28 (Fig. S1B-C). *Mllt11* transcript levels decreased but remained detectable in the hippocampus on through P28 (Fig. S1C).

Given the temporal and spatial overlap between *Mllt11* expression and neurogenic regions of the cerebrum, we set out to investigate the role of Mllt11 in the development cerebral cortex, and specifically in the formation of the upper layer (UL) 2/3 callosally projecting CPNs that bridge communication interhemispherically (Fame et al., 2011) where *Mllt11* expression appeared highest during corticogenesis (Fig. S4A). We utilized the targeted *Mllt11* null allele to generate a conditional knockout (cKO) Mllt11 allele in which the entire coding sequence in Exon 2 is flanked by loxP sites. To avoid non-specific effects of the selection cassette, mice encoding germline expressing flp recombinase were crossed to mice harbouring the *Mllt11* targeted allele, leaving only loxP sites used to excise the entire protein-coding exon to create a conditional allele. Subsequently, in order to inactivate the *Mllt11* in UL2/3 of the cerebral cortex, we crossed the *Mllt11^Flox/Flox^* mice to the *Cux2iresCre* strain along with the *Ai9 TdTomato* reporter mice to visualize recombined neurons, creating *Mllt11* conditional knockout mutants (cKO). Cux2 is highly expressed in developing UL2/3 projections neurons, and mosaically expressed in the SVZ in the pallial cortex (Cubelos et al., 2008a; Cubelos et al., 2008b; Nieto et al., 2004; Zimmer et al., 2004). The resulting *Cux2iresCre*-driven *Mllt11* cKO mice should lack *Mllt11* expression in UL of the neonatal cortex. We confirmed *Mllt11* loss in UL CPNs qPCR of E18.5 cortices (Fig. S2A-B) as well as by ISH at P7 (Fig. S2C-D). Due to our genetic strategy, all cells exhibiting *Mllt11* loss were labeled by TdTomato fluorescence. TdTomato labeling was observed mostly in UL 2/3 of the cortex (Fig. S3A) and commissural tracts at P7 (cc; Fig. S3B), confirming that Cre-medicated excision was specific to the *Cux2* expression domain.

### Loss of *Mllt11* from Upper Layer 2/3 Leads to Progressive Decrease in Cortical Thickness

Brain and body weight were not found to differ significantly between *Mllt11* cKO mutants and controls at E18.5, though there was a very slight trend toward decreased brain weight in the cKOs (Fig. S4B-D). Mutants displayed a thinning of the cortex visible in the cortical plate (CP) and white matter (WM) beneath it (Fig. S4E; Fig. 4A, D). To explore the origin of this cortical thinning, we examined the morphology of the cortex beginning at E14.5 (Fig. S4F) when UL CPNs are born and begin to express Cux2 (Cubelos et al., 2008a; Zimmer et al., 2004). The thinning of the cortex was observed beginning at E16.5 (Fig. S4G), increasing in severity by E18.5 (Fig. S4H). We noted a non-significant trend toward fewer total DAPI+ cells in the *Mllt11* mutant cortex (Fig.S4H). However, we report significant changes in the distribution of DAPI+ nuclei in deeper bins of the *Mllt11* cKOs, with a larger proportion of DAPI+ nuclei localized apically in E18.5 mutants relative to controls (Fig. S4F-H). We also evaluated whether programmed cell death could account for the cortical thinning, but did not find any cells positive for the apoptotic marker cleaved caspase 3 (CC3) in the *Mllt11* neonatal cortex (Fig S5A-B), while retrosplenial regions, which typically show enhanced apoptosis, showed comparable levels of CC3+ staining in controls and mutants (Fig. S5C).

*Mllt11* expression was restricted to the developing cortical plate and not in the apical progenitor regions (VZ) of the cortex (Fig. S4A). To rule out potential cell non-autonomous contributions from neural progenitors to the *Mllt11* mutant cortical phenotype we quantified cells positive for progenitor markers from E14.5-18.5, corresponding to the period of UL neurogenesis. Pax6, a radial glial cell (RGC) marker (Gotz et al., 1998), was unaltered at all observed time points (Fig. S6A-C). Sox2, a less restrictive marker for neural progenitors and quiescent RGCs (D’Amour and Gage, 2003; Ellis et al., 2004), also exhibited no significant differences in expression (Fig. S6D-F). Tbr2+ intermediate (basal) progenitors, which give rise to UL 2/3 CPNs, did not differ markedly between control and *Mllt11* cKO groups, but displayed fewer cells within deep (non-progenitor) regions of the mutant cortex at E18.5, suggesting alterations in the formation and/or migration of nascent neurons from the SVZ (Fig. S6G-I).

### *Mllt11* is Required for the Maintenance of UL CPN Molecular Identity

The transcriptional de-repression loop that specifies cell types in discrete cortical laminae coincides with the birth and migration of projection neurons (Kumamoto et al., 2013; Toma et al., 2014). Thus a loss of *Mllt11* in UL progenitors can potentially affect neuronal birth, migration, and/or specification. We therefore investigated whether *Mllt11* loss had any role in regulating the molecular identity of UL CPNs. Expression of Satb2 in CPNs of L2-4 exhibited a decrease at E18.5 (Fig. 1A) as did CDP/Cux1, another marker of UL CPNs (Fig. 1B). Investigation at earlier developmental time points revealed that this phenotype was progressive, beginning at E16.5 and increasing in severity by E18.5 (Fig. S7A-D). While the extent of CDP/Cux1 and Satb2 staining displayed variability across individual mutants, there was a consistent decrease in expression levels and downward shift in expression domain in all *Mllt11* cKOs. TdTomato labelling was used to identify Cux2+ cell labeling in intermediate progenitors that give rise to UL2/3 CPNs, and while reporter levels were found to be unaltered at all time points, there was a consistent downward shift in expression domain at E18.5 (Fig, 1C, Fig. S7E-F). Collectively, these findings suggested that *Mllt11* loss in nascent UL neurons did not affect their neurogenesis, or the activation of UL gene expression programs, but may have affected their migration to contribute to L2/3 formation.

**Figure 1:**
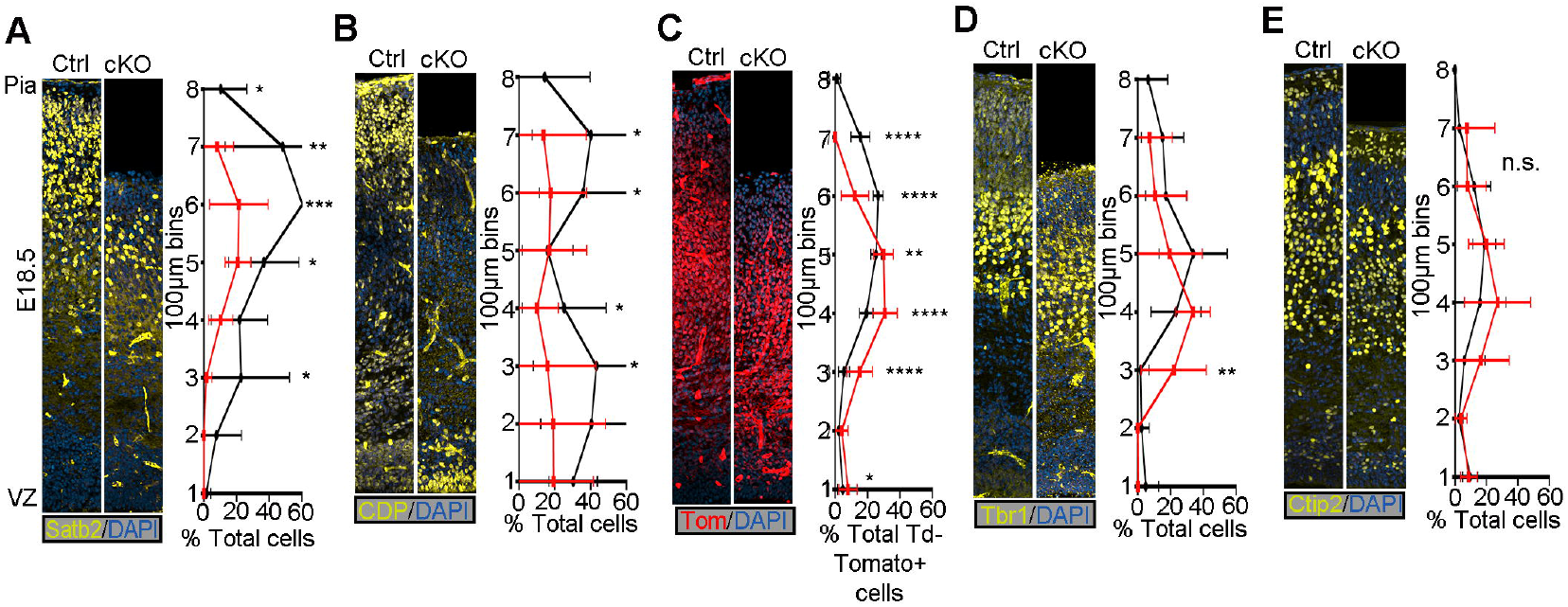
*Mllt11* loss perturbed formation of UL CPNs without affecting deeper layers. (A-E) Coronal cortical slices at E18.5 stained for cortical layer markers with IHC. (A) *Mllt11* conditional knockout (cKO) cortical slices stained for Satb2 displayed a severe decrease in expression at E18.5 compared to controls (ctrl). (B) CDP/Cux1 levels were decreased in *Mllt11* cKOs compared to controls at E18.5. (C) Distribution of TdTomato expression in the cortex at E18.5 was comparable between controls and cKOs. (D-E) Tbr1 (D) and Ctip2 (E) levels were comparable between controls and cKOs but their expression domains exhibited a downward shift at E18.5. Line charts represent percentage of positive cells normalized to DAPI+ nuclei per 100μm x 100μm bin for (A-B) and (D-E) and percentage of total TdTomato+ cells per 100μm x 100μm bin for (C). Welch’s t-test, (A, D, E) N=4, (B-C) N=5. Data presented as mean ± SD. n.s., not significant. *P≤0.05; **P≤0.01; ***P≤0.001; ****P≤0.0001. Scale bar: 50μm for (A-O). VZ, ventricular zone.

The cortical de-repression loop functions on a cellular level by alternately repressing DL and UL fate (Toma et al., 2014) so we wanted to determine if the loss of UL CPN fate was coupled with a complementary upregulation of DL CPN fates in these cells. We examined the localization of DL6- and DL5-specific markers Tbr1 (Fig. 1D) and Ctip2 (Fig. 1E) respectively, which both exhibited similar numbers and staining patterns between *Mllt11* cKOs and controls at all time points (Fig. S7G-J). However, the mutants displayed a downward shift of the expression domains of both markers consistent with the decreased cortical thickness and UL formation (Fig. 1D-E). To determine whether the loss of CPN subtype-specific markers was indicative of loss of neuronal identity, we examined the localization of NeuN, a marker for maturing neurons (Mullen et al., 1992) and found no alterations in *Mllt11* cKOs (Fig. S8A-B). No changes in GFAP staining, a marker of radial glial fibres and astrocytes (Bignami and Dahl, 1977; Zhang, 2001) was apparent, demonstrating that these *Mllt11* mutant cortical cells were not experiencing a shift from neuronal to glial identity (Fig. S8C). This is consistent with the neuronally restricted expression of Mllt11 (Fig. S1C, S4A) (Yamada et al., 2014).

Given that the *Mllt11* mutants failed to maintain UL-specific transcription factors over time, we wanted to understand whether the cortical de-repression loop and layer-specific celltype specification were de-coupled, such that CPNs born at the time of UL CPN birth could instead mis-express a DL-specific transcriptional regulator. Pregnant dams were injected with EdU at E14 or E16 to label UL neurogenesis and embryos were harvested at E18.5 to examine the co-localization of the DL CPN marker Ctip2 with EdU. The proportion of Ctip2+ CPNs born at E14 did not differ significantly between controls and cKOs with the exception of the overall downward shift in the distribution of Ctip2+/EdU+ cells in the cKOs (Fig. S9A-a’). Embryos pulsed with EdU at E16 and harvested at E18.5 displayed the same trend, exhibiting a downward shift but no significant differences in overall proportions of Ctip2+/EdU+ co-labeled cells (Fig. S9B-b’). Taken together, these data suggest that UL CPNs are not experiencing a fate change to DL CPNs, but initiate expression of proper layer-specific transcription factors which they fail to maintain over time.

### *Mllt11* is Required for the Migration of CPN Progenitors

Given the thinning of the cortex and downward shift of cortical layer markers throughout development, we hypothesized that Mllt11 may regulate CPN migration into the CP. To explore this, we pulsed pregnant dams with EdU to label UL CPNs at E14 or E16 and observed the localization of EdU at E18.5. CPNs labeled with EdU at E14 exhibited a severe downward shift of roughly 100 μm, consistent with the decrease in cortical thickness (Fig. 2A). A greater proportion of E14-born CPNs remained restricted to the deeper bins of *Mllt11* cKO cortices relative to controls (Fig. 2A). As CPNs labeled with EdU at E16 were still in transit by E18, with only a few control CPNs reaching the outermost layers of the cortex, the discrepancy in UL cell migration was less pronounced in cKOs (Fig. 2B). To examine whether *Mllt11* loss affected the neurogenesis of UL neurons, we pulsed dams with EdU for 2 hours at E14, which efficiently labeled nuclei within the basal progenitor region (Fig. 2C). We observed no difference in the numbers or distribution of EdU+ nuclei in *Mllt11* mutants vs. control cortices after a short pulse at E14 (Fig. 2C). We repeated this short pulse at E18, when UL neurogenesis is nearly complete and noted no differences in EdU+ cells in the superficial cortex of *Mllt11* mutant vs. control brains (Fig. 2D). We did observe a slight increase of EdU+ cells in the deepest 100 μm bin of the *Mllt11* mutant cortex relative to controls, possibly reflecting aberrant migration of later born neurons (Fig. 2D). Overall, the EdU birth dating analysis suggested that Mllt11 primarily regulates the migration, but not neurogenesis, of UL CPNs into the cortical plate.

**Figure 2:**
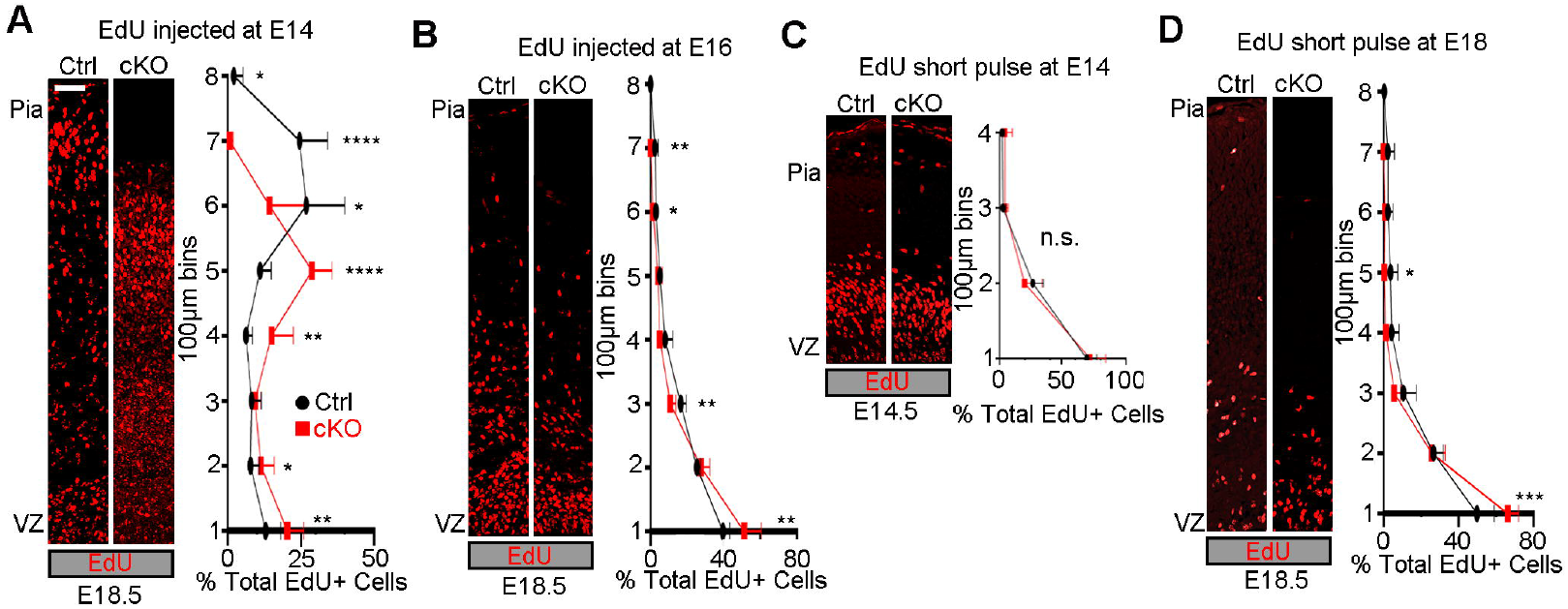
EdU birthdating demonstrating a CPN migratory defect in *Mllt11* cKO mutants. (A-B) EdU birthdating of UL CPNs by injection at E14 showed altered distribution in cKOs at E18.5, reflecting a downward shift in expression correlating with the decreased cortical thickness. (B) Moderate downward shift in migrating CPNs in cKOs relative to controls following an EdU pulse at E16 (B). (C) Proliferating cells labeled with a short pulse of EdU at E14 showed no significant differences in numbers or distribution between controls and cKOs. (D) No significant difference in numbers of nuclei labeled by a short pulse of EdU at E18 populating most bins in cKOs vs. controls, but a greater proportion of cells were retained in the cKO VZ region. Line charts represent percentage of positive cells per 100μm x 100μm bin as a proportion of total EdU+ cells. Welch’s t-test, (A, C) N=4, (B, D) N=3. Data presented as mean ± SD. n.s, not significant. *P≤0.05; **P≤0.01; ***P≤0.001; ****P≤0.0001. Scale bar: 50μm for (A-O). VZ, ventricular zone.

A more conclusive test for a role of Mllt11 in the migration of CPN progenitors into the cortical plate would be to overexpress it in the developing fetal brains. We therefore used *in utero* electroporation to express a bicistronic vector containing *Mllt11* and *GFP* cDNAs, or control *GFP* vector alone, in wild type mouse embryonic forebrains. We performed electroporations in anesthetised pregnant dams at E13.5, by injecting a DNA solution containing expression vectors unilaterally into forebrain ventricles, and analyzed the resulting fetal brains 2 days after at E15.5, when neurons are migrating into the cortical plate. At E15.5, CPNs overexpressing *Mllt11* primarily colocalized with preplate marker Tbr1 (Fig. 3A), while GFP-expressing control CPNs localized more closely with Tbr2+ basal progenitors near the IZ (Fig. 3B-D), demonstrating *Mllt11* overexpression is sufficient to promote CPN migration into the developing cortical plate. Taken together, our loss and gain-of-function analyses strongly support a role for Mllt11 in promoting nascent UL neuronal migration in the developing cortical plate.

**Figure 3:**
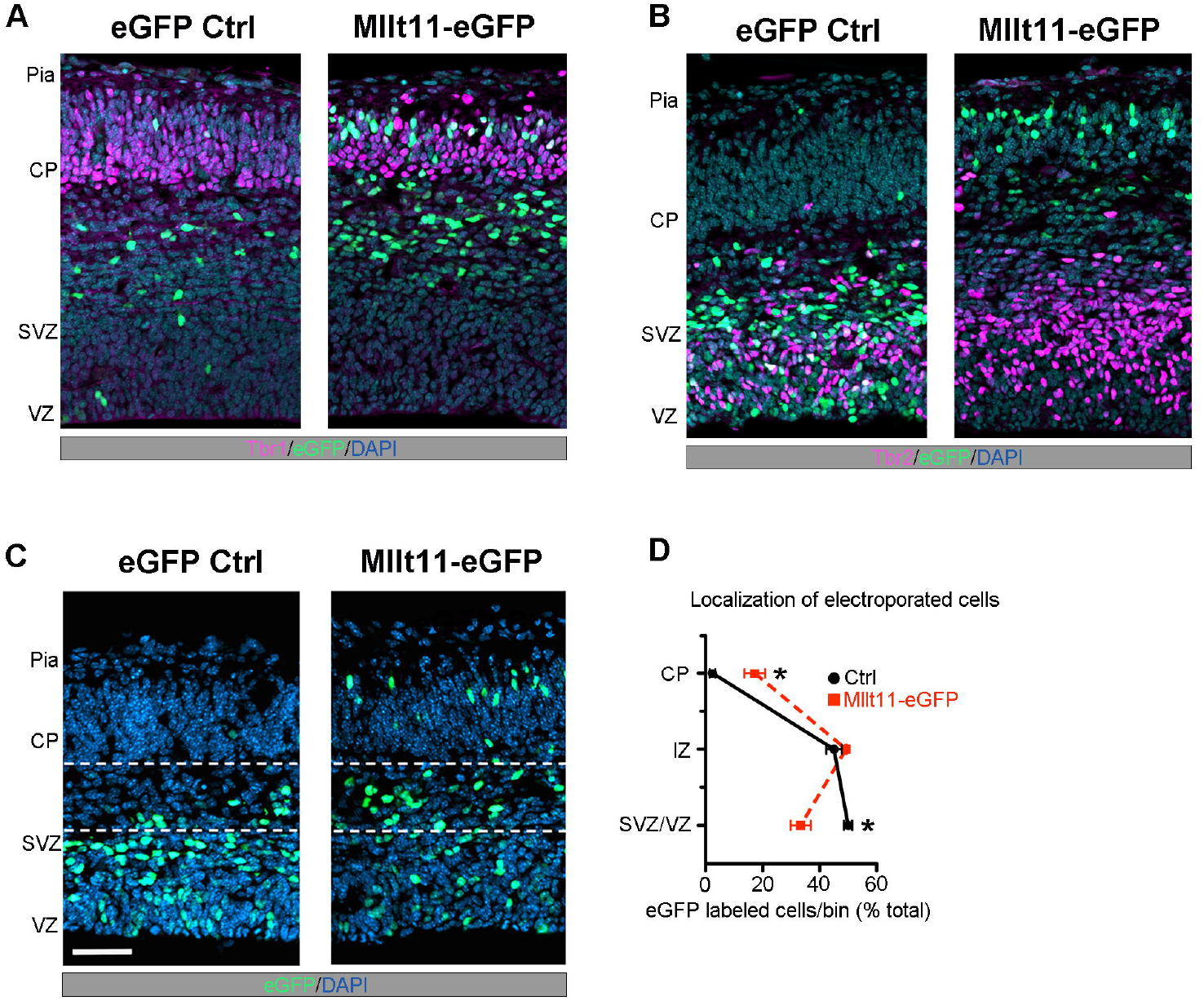
*Mllt11* overexpression promoted migration into the cortical plate. (A-B) E15.5 coronal cortical sections after electroporation at E13.5 wither either a control eGFP (left) or *Mllt11-ires-eGFP* (Mllt11-eGFP) bicistronic plasmid (right). (B) *Mllt11-eGFP* electroporation promoted migration into the CP, identified by Tbr1 staining. (C) Control GFP (eGFP-control) electroporated cells remained mostly within the SVZ/VZ, identified by Tbr2 staining. (D-E) Localization of eGFP-control and Mllt11-eGFP+ cells (D) quantified by cortical region (E). Welch’s t-test; N=3 Mllt11-eGFP, N=4 eGFP controls. Data presented as mean ± SD. *P≤0.05. Scale bar: 25μm for (A). CP, cortical plate; IZ intermediate zone; SVZ, subventricular zone; VZ, ventricle zone.

The migration of CPN progenitors is facilitated by a meshwork of glial fibres which extend from the VZ towards the pial surface (Campbell and Gotz, 2002; Hartfuss et al., 2001; Miyata et al., 2001). These fibres act as tracks for extending neurites guiding neuronal translocation, contributing to the meshwork laid down by earlier born progenitors. As such, the orientation of Nestin+ fibres can indicate the direction of migration into the CP (Gotz et al., 1998; Nadarajah et al., 2001). We wanted to rule out any possible cell non-autonomous effects of *Mllt11* loss on the migrating scaffold in the developing cortex by carefully analyzing the Nestin+ glial scaffold of the cortex. We found no significant alterations in orientation, density, and dispersion of RGC fibres in *Mllt11* cKOs during cortical development (Fig. S10), but did note a non-significant trend toward a greater dispersion of the fibres in cKOs relative to controls at E18.5 (Fig. S10L). Guidance of migratory CPNs is dependent on the earliest born Cajal-Rietzus (CR) cells of L1, which secrete Reelin to attract and halt neuronal migration at the pial interface (Hirota and Nakajima, 2017; Tueting et al., 1999). However, there were no differences in the expression of CR markers p73 or Reelin at E14.5 (Fig. S11A) or E18.5 (Fig. S11B) following *Mllt11* loss in UL progenitors and nascent neurons. Altogether, these findings demonstrate that the aberrant migration of UL CPNs in *Mllt11* mutants was not caused by alterations in the development of Reelin-secreting CR cells, nor abnormalities in radial glial fibre scaffold. Rather, Mllt11 seems to be required cell autonomously to promote invasiveness of nascent neurons into the developing cortical plate.

### *Mllt11* Loss Leads to Reduced Callosal Crossing Fibres and Thinning of WM tracts

The migratory defects alone may be insufficient to explain the decreased cortical thickness of *Mllt11* cKO brains at E18.5 (Fig.S4). Given the reduction in UL CPNs (Fig. 1) would largely affect interhemispheric projections (Aboitiz and Montiel, 2003; Alcamo et al., 2008; Fame et al., 2011), we next examined whether *Mllt11* loss impacted formation of projection fibres contributing to the cortical white matter (WM) tracts by staining for NF in coronal brain slices. We observed notable decreases in the thickness of NF+ fibres in the WM tract and in the proportion of the cortex containing WM between E16.5 (control=235 μm ± 49.68, cKO=172 μm ± 17.06; Fig. 4A-C) and E18.5 (control=235.1 μm ± 12.58, cKO=200.8 μm ± 15.66; Fig. 4D-F) following *Mllt11* loss. Importantly, while NF+ fibre staining in the corpus callosum was greatly reduced in *Mllt11* cKOs at E18.5, NF+ fibres in the internal capsule, reflecting corticothalamic projections, remained unchanged in the mutant brains (ic; Fig. 4G). Taken together, our findings revealed a critical role for Mllt11 in UL CPN neurite morphology and axonogenesis of callosal fibres connecting the developing cerebral hemispheres.

**Figure 4:**
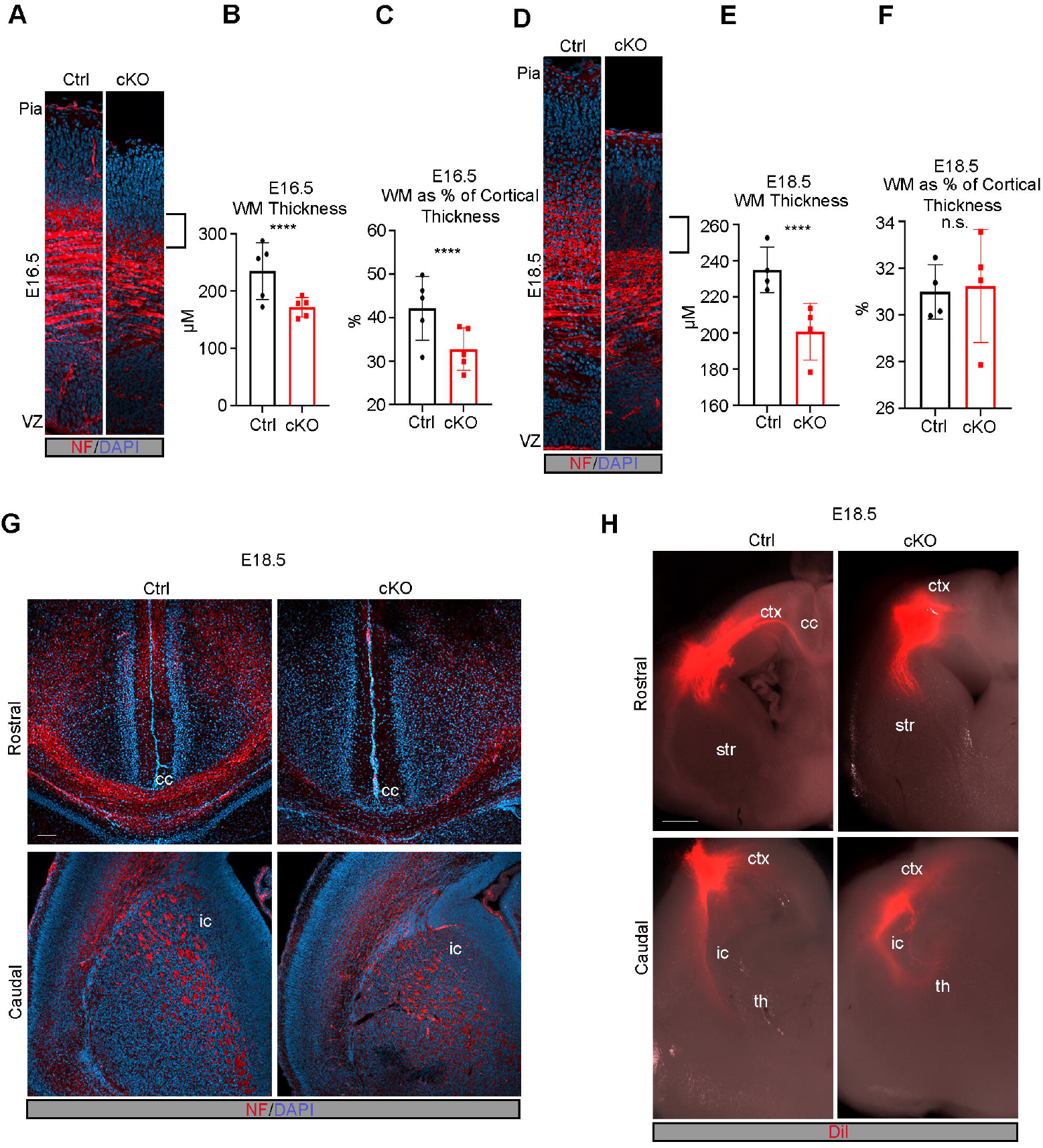
Formation of white matter tracts and callosal projections is impaired in the *Mllt11* cKO cortex. (A) Image of cortical white matter tracts labeled with neurofilament at E16.5. Brackets indicated decreased white matter. (B-C) Quantification of cortical white matter thickness (B) and white matter thickness as a proportion of total cortical thickness (C) showed significant decrease in white matter in cKOs relative to controls at E16.5. (D) Image of cortical white matter tracts labeled with neurofilament at E18.5. Brackets show decreased white matter. (E-F) Quantification of cortical white matter thickness (E) and white matter thickness as a proportion of total cortical thickness (F) displayed a decrease in cKOs at E18.5. (G-H) Coronal sections of E18.5 cortices at rostral (upper panels) and caudal (lower panels) axial levels labeled with neurofilament (G) or traced with DiI (H). (G) Neurofilament labeling of the corpus callosum was significantly decreased in cKO compared to controls but labeling of the internal capsule was unaffected. (H) DiI labeling was absent in the corpus callosum of cKO slices while control cortices displayed crossing fibres labeled by DiI. Rostrally, the internal capsule was traced comparably in control and cKO cortices. Welch’s t-test, (A-C) N=5, (D-F) N=4, (H) N=3. Data presented as mean ± SD. ****P≤0.0001. Scale bar: 200μm for (A); 50μm for (B). ctx, cortex; cc, corpus callosum; str, striatum; th, thalamus; ic, internal capsule; VZ, ventricular zone; WM, white matter.

The reduction of NF+ fibres crossing the corpus callosum in mutant brains suggested that *Mllt11* loss impacted the ability of UL2/3 CPNs to extend fibres across the telencephalic midline. To address this, we injected the lipophilic fluorescent dye DiI into the WM of cortical slices at E18.5 and allowed it to diffuse through the tissue for 2 weeks before visualization. DiI labeling was not detectable in the callosal fibres of *Mllt11* cKOs, while controls exhibited clearly labeled crossing fibres (Fig. 4H). In contrast, the corticothalamic projections of the internal capsule were intact and traced by the DiI in both controls and cKOs, demonstrating that corticothalamic projections from DL 6 of the cortex were unaffected by the *Cux2iresCre* driven *Mllt11cKO* strategy (ic; Fig. 4H). Altogether, our findings demonstrated that *Mllt11* loss in UL neurons specifically impacted the development of callosal projections.

### Mllt11 Interacts with Cytoskeletal Proteins

Our birthdating analysis suggested that the migratory defects observed in *Mllt11* cKOs are likely independent of the maintenance of UL transcriptional programs. To identify potential pathways through which Mllt11 might be exerting its effects on CPN migration, we performed a GST pulldown assay to identify interacting proteins using lysate from E18.5 brains. Potential Mllt11-interacting proteins are listed in Table 1 and include multiple α and β tubulin isoforms (> 70% coverage) as well several atypical myosins, such as Myosin 5a (36%), Myosin 9 (24%) and Myosin 10. A potential association between Mllt11 and tubulin is consistent with a recent whole cell proteomic study, which identified Mllt11 as a likely interactor of both α- and ß-tubulins (Go et al., 2021). Given that acetylation of α-tubulin is associated with stabilized microtubules and is crucial for outgrowth of stable neurites, we decided to focus on validating interactions between Mllt11and acetylated α-tubulin by immunoprecitiation analysis. We overexpressed a myc-tagged Mllt11 in Hek293 cells and pulled down acetylated-α-tubulin (Fig. 5A). Probing the blots with anti-Mllt11 antibodies confirmed this interaction was dependent on heterologous Mllt11 expression in the cells (Fig. 5B). We also confirmed an association between Mllt11 and Myosin 10 (data not shown). Thus, we validated an association between Mllt11 and stabilized tubulin isoforms.

**Table 1.**
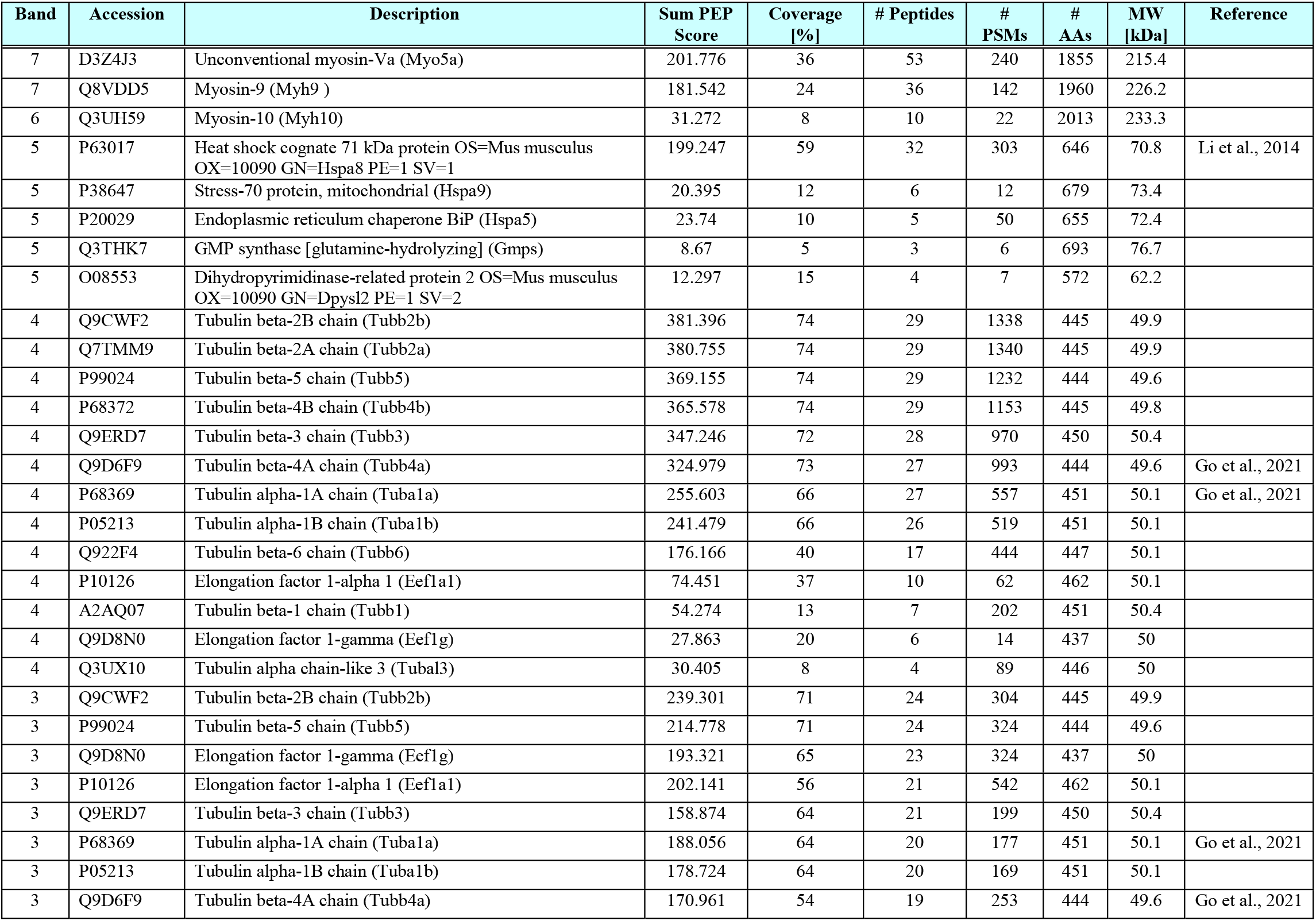
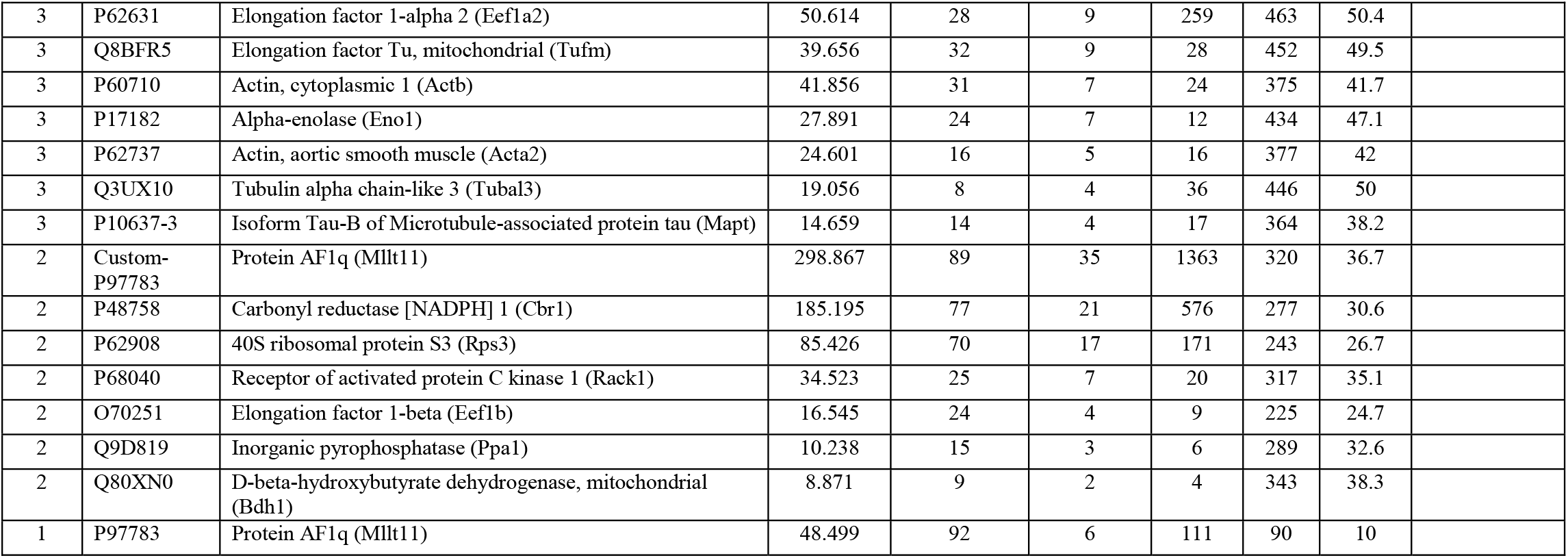
Potential Mllt11 interaction targets from GST pull downs in E18.5 whole brain lysates.

**Figure 5:**
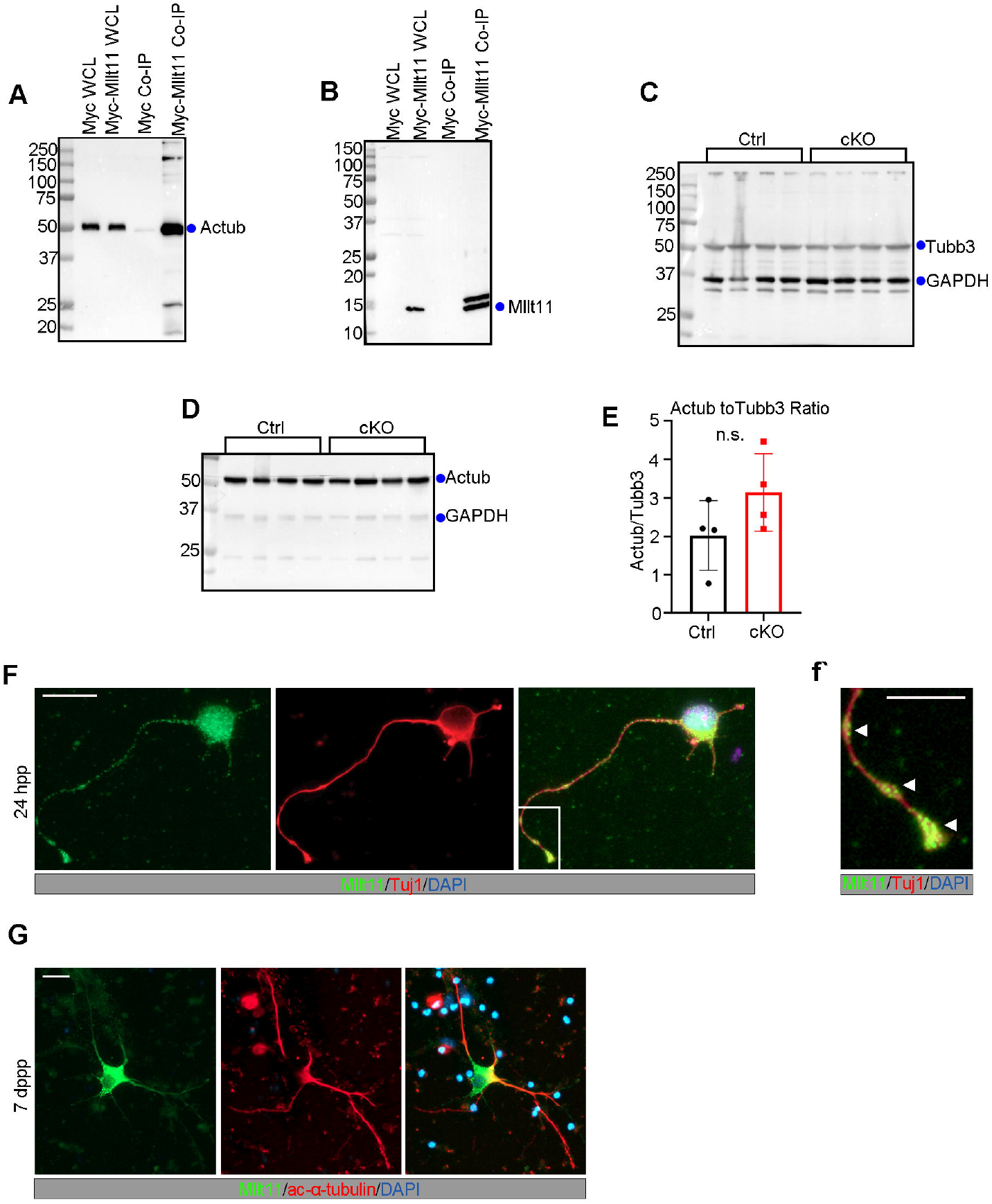
Mllt11 associates and colocalizes with acetylated α-tubulin in growing neurites. (A) Co-immunoprecipitation of acetylated α-tubulin (Actub) with Myc-tagged Mllt11 in whole cell lysates of HEK293 cells compared to a myc programmed control. (B) Co-IP probed with Mllt11 antibody showing IP signal restricted to lanes containing Mllt11. (C-D) Western blots of E18.5 brain lysates probed for Tubb3 (C) and acetylated α-tubulin (D) show no differences in levels between controls and cKOs. (E) Bar graph of the ratio of acetylated tubulin to Tubb3 normalized to GAPDH. Conditional mutants trend toward an increased ratio (not significant). (F-G) Primary cortical neurons cultured for 24 hours (F) or 7 days (G) post-plating. (F) After 24 hours, Mllt11 colocalizes with Tubb3 in the growth cone and at swellings along the distal axon as indicated by arrowheads (f’), and in the soma (F). After 7 days, Mllt11 was primarily found along proximal portions of the axon where it colocalizes with acetylated α-tubulin (G). Welch’s t-test, N=4 (controls and cKOs). Data presented as mean ± SD. n.s, not significant (i.e. P>0.05). Scale bar: 20μm.

To determine whether Mllt11 regulates tubulin expression, western blots were performed on lysates from *Mllt11* cKO and control E18.5 cortices. Tubb3 (a β-tubulin isoform) and acetylated α-tubulin levels appeared comparable between E18.5 *Mllt11* mutant and controls cortices (N=3, Fig. 5C-D). Interestingly, there was a trend towards an increase in the ratio of acetylated tubulin to Tubb3 in *Mllt11* cKOs, although it was not statistically significant (Fig. 5E). This suggested the possibility that *Mllt11* loss affected the formation of stabilized microtubules.

To evaluate whether Mllt11 localized to growing neurites, we used immunocytochemistry to probe the sub-cellular distribution of Mllt11, Tubb3, and acetylated α-tubulin in cultures of primary fetal cortical neurons. Cortical neurons were extracted from wild type embryos at E18.5 and cultured for 24 hours or 1 week, then immunostained to reveal the localization of Tubb3, acetylated α-tubulin, and Mllt11. After 24 hours *in vitro*, corresponding to the extension of a primary neurite, Mllt11/Tubb3 co-localized in discrete punctate patterns in varicosities along the distal portion of the developing neurite, as well as in the growth cone (Fig 5F-f’). Limited overlap was seen in the soma (Fig. 5F). After 1 week in culture, neurons displayed a more elaborate neurite arborisation pattern and acetylated α-tubulin and Mllt11 were both detected along proximal portion of neurite (Fig. 5G). In summary, Mllt11 expression displayed a high degree of overlap with acetylated α- and β-tubulin isoforms differentially over neurite development, consistent with a role in neurite development. This is in agreement with a study identifying possible microtubule interactions with Mllt11 (Go et al., 2021)

### Mllt11 is Required for Neurite Outgrowth and Extension

The relationship between stabilized and dynamic forms of tubulin has been shown to regulate the migratory potential of neurons and neurite extension due to the cycling of microtubule severing during outgrowth (Lin and Smith, 2015; Sudo and Baas, 2010; Wei et al., 2018). If *Mllt11* loss affected the relative amounts of stabilized microtubules, we would expect to see significant changes in neurite morphogenesis of UL neurons. We therefore evaluated the morphology of cultured primary cortical neurons from *Mllt11* cKOs. We took advantage of the Ai9 tdTomato+ reporter introduced into the *Cux2iresCre/+; Mllt11^Flox/+^* conditional mouse to efficiently label and isolate UL primary neurons in culture. After 24 hours *in vitro*, the length of MAP2+ primary neurites was reduced in cKOs compared to controls (control=83.78 μm ± 23.17, cKO=73.62 μm ± 24.52, N=150; Fig. 6A-B), as was the total length of all neurites (control=127.4 μm ± 56.11, cKO=92.34 μm ± 28.15, N=150; Fig. 6C). Moreover, the proportion of total length of extended neurites represented by the primary neurite at 24 hours was slightly increased in cKOs (control=.72 ± .21, cKO=.81 ± .18, N=150; Fig. 6D). However, after 1 week of culture *in vitro*, the total neurite length was drastically reduced in *Mllt11* cKO CPNs (control=1415 μm ± 1112, cKO=595.9 μm ± 597.2, N=150; Fig. 6E). Since *Mllt11* loss progressively attenuated primary neurite outgrowth in cultured primary UL CPNs, we next evaluated a requirement for Mllt11 in the elaboration of dendritic arborisation patterns in cultured CPNs. An advantage of using cultured neurons is that they are free of apical and basal contacts, allowing their neurites to radiate uniformly from the soma. This allows for the evaluation of dendritic complexity by Sholl analysis. *Mllt11* mutant primary neurons displayed much less complexity in the branching of their neurites (Fig. 6F-G), thus *Mllt11* loss greatly impacted the arborisation patterns of cultured UL CPNs.

**Figure 6:**
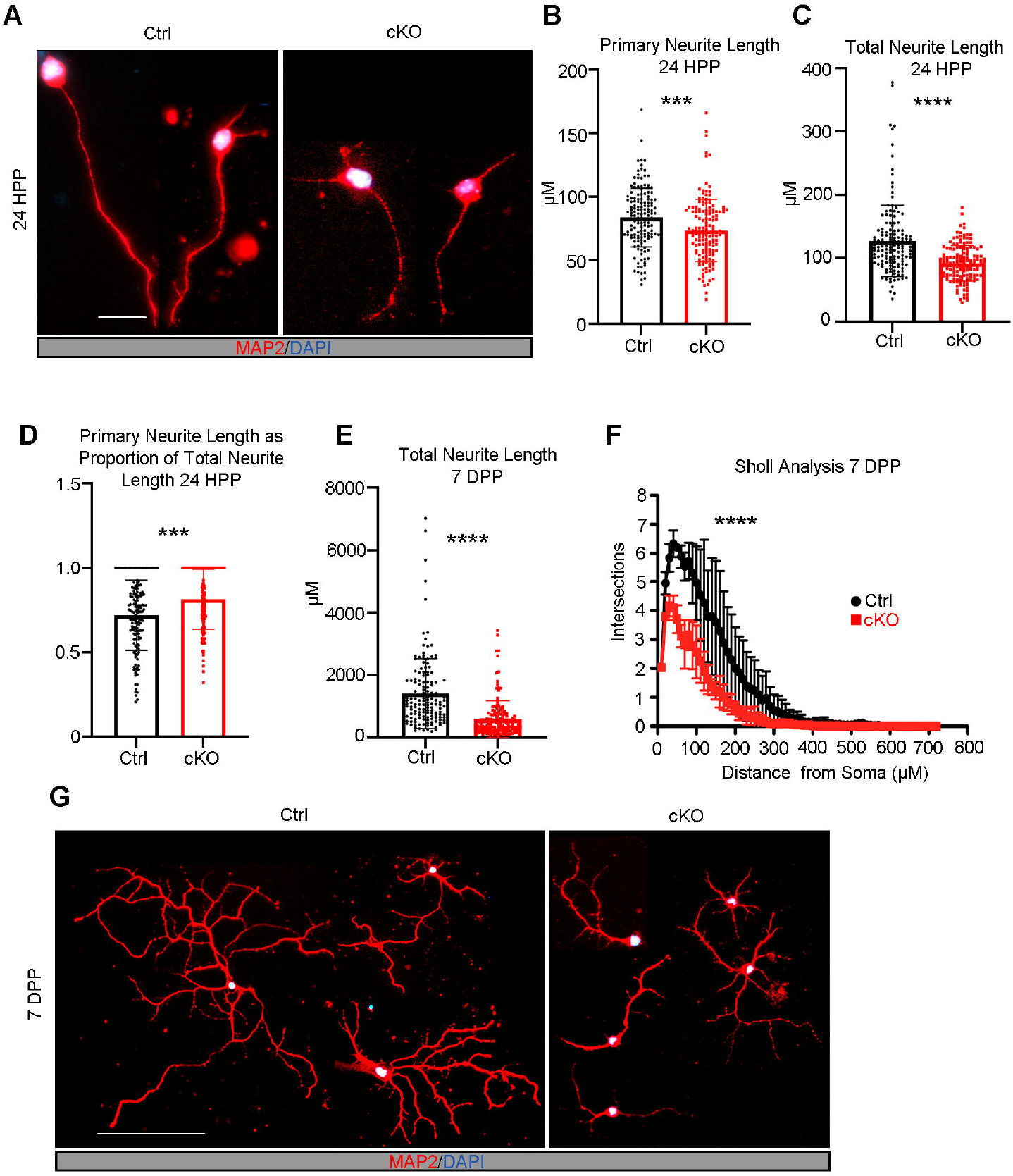
*Mllt11* loss decreased neurite outgrowth and branching complexity *in vitro*. (A) Primary cortical neurons derived form E18.5 brains cultured for 24 hours post-plating. Neurites identified with MAP2 staining (red). (B-C) 24 hours post-plating, primary neurite length (B) and total neurite length (C) were significantly decreased in cKO neurons relative to controls (D). The proportion of total primary neurite length was significantly increased in cKO neurons relative to controls. (E-G) Quantification of decreases in total neurite length (E) and branching complexity (F) of cKO relative to control primary cortical neurons cultured for 7 days post-plating (G). Paired t-test, for both controls and cKOs N=150 neurons (50 neurons/individual X 3 individuals). Data presented as mean ± SD. *P≤0.05; **P≤0.01; ***P≤0.001, ****P≤0.0001. Scale bar: 20μm for (A); 50μm for (E). HPP, hours post-plating; DPP, days post-plating.

Given the vast array of morphological growth and pruning in early perinatal development, we next evaluated the requirement of Mllt11 in the formation of dendritic arborisation morphology characteristic of UL CNs *in vivo* at a time when mature arborisation patterns have fully developed. Golgi staining was performed on *Mllt11* cKO and control brains harvested at P28 to capture mature CPN morphologies (Fig. 7). Importantly, we observed a severe decrease total neurite length in UL2/3 CPNs in cKOs relative to controls (control=1176.62 μm ±164.73, cKO=563.85 μm ±106.1, Fig. 7A-D). The extent of decrease of neurite aborisation *in vivo* was similar to that observed in cultured primary neurons. Sholl analysis revealed that the overall neuronal morphology was also much less complex with less elaborately-branched neurites in *Mllt11* cKOs (Fig. 7E). Taken together, these findings confirmed a critical requirement of Mllt11 in the growth and elaboration of neurites during the development of mature superficial CPN morphologies and projection phenotypes.

**Figure 7:**
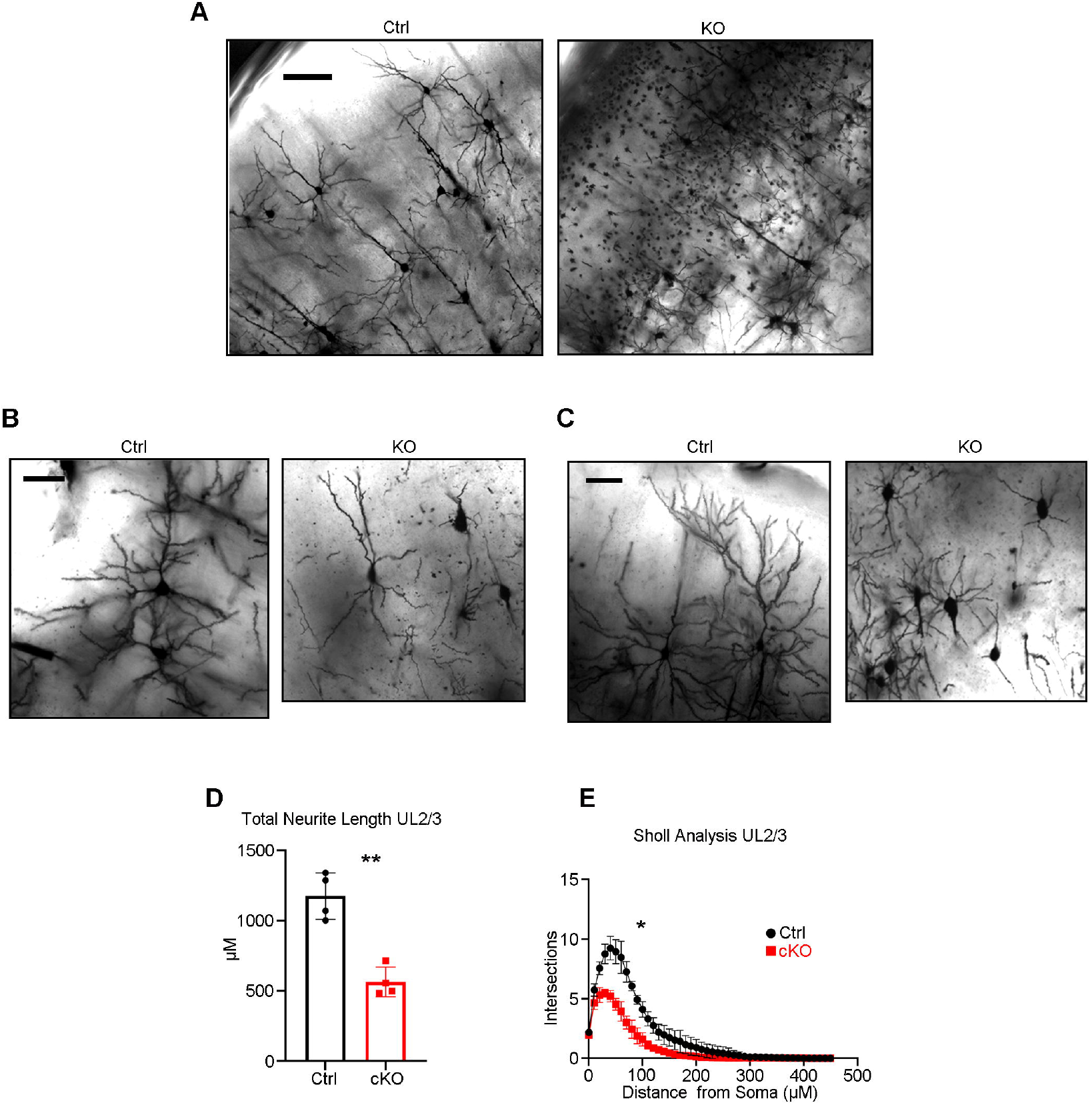
Decreased neurite outgrowth and branching complexity in UL CPNs of the *Mllt11* cKO cortex. (A-C) Corresponding images from coronal sections of P28 Golgi stained control and cKO cortices. (A) 10x magnification view of overall cortical morphology. cKOs displayed dark, densely stained punctate structures in the upper cortical layers, which were absent in controls. (B-C) 200x magnification view of sample UL2/3 CPNs varying in morphology between control and cKO neurons. (D-E) Quantification of neurite length (D) and branching complexity (E) within UL2/3 showed reduced neurite length and branching complexity in cKOs compared to controls. Length measurements analyzed with Welch’s t-test, layer-specific Sholl analysis compared with paired t-test, N=4 (controls and cKOs), 30 neurons/individual. Data presented as mean ± SD. *P≤0.05; **P≤0.01. Scale bar: 100μm for (A); 50μm for (B-C).

## DISCUSSION

We described the role of a novel type of a vertebrate-specific neuronal restricted protein, Mllt11, which acts as a regulator of neurite extension and migration of superficial cortical projection neurons. Using the *Cux2iresCre* driver to excise *Mllt11* in most developing UL cortical neurons, we revealed that *Mllt11* loss led to decreased cortical white matter tracts and callosal fibres crossing the forebrain midline, as well as an inability to extend and maintain mature, arborescent dendrites. Nestin+ and GFAP+ fibres and progenitor markers were largely intact in the absence of *Mllt11*, consistent with the expression of *Mllt11* in the developing cortical plate, but not in VZ progenitors. Furthermore, *Mllt11* conditional mutants did not affect the formation of layer 1 Reelin+ cells, which guides the migration of cortical neurons and formation of fibre scaffolds, suggesting instead that *Mllt11* loss acted cell autonomously to affect the radial migration of CPN progenitors into the cortical plate. The inefficient migration of UL CPNs in *Mllt11* mutant brains likely reflected a role for Mllt11 in regulating the cytoskeleton, as is associated efficiently with stabilized microtubules. Neurons lacking Mllt11 displayed deficits in development of arborescent neurites both *in vivo* and in *vitro*. This neurite extension deficit in the mutants was further confirmed by the decrease in neurofilament and DiI staining of callosal fibres, and thinning of white matter tracts of the cortex, demonstrating that Mllt11 is a regulator of neurite outgrowth and maintenance required for forebrain connectivity.

Cortical thinning and increased ventricular lumen surface area are common phenotypes associated with neurodevelopmental disorders such as autism, suggesting cytoskeletal dysregulation may underlie the occurrence of a subset of these disorders (Bahi-Buisson et al., 2014; Li et al., 2019; Mensen et al., 2017). We now show that Mllt11 interacts with tubulin isoforms, confirming findings by others in a recent proteomic screen (Go et al., 2021), which provide insight into the mechanistic basis of the *Mllt11* mutant phenotype. The microtubule cytoskeleton generates polarized force and provides a functional railway along which motor proteins can migrate, whether the function be to drive actin polymerization in generating protrusive forces at the leading edge of the growth cone or integration of extracellular cues into regulation of microtubule activity (Buck and Zheng, 2002; Dent and Kalil, 2001; Lee et al., 2004; Zhou et al., 2004). Tubulin mutations have been shown to impair neuronal migration into the cortex (Aiken et al., 2017), and can specifically affect UL CPNs (Keays et al., 2007), which rely on a more dynamic, multipolar mode of cytoskeletal reorganization as they extend short processes to “crawl” through established DL CPNs (Miyata et al., 2001; Sakakibara et al., 2014). Mutations in α-tubulin specifically have been shown to cause axonal trafficking defects that impair synaptic stability and function without impacting axonal degeneration or neuronal survival (Buscaglia et al., 2020). Potential interactions of Mllt11 with myosins (data not shown) and microtubules (this study) could provide a mechanistic explanation for the impaired migration and morphology seen in *Mllt11* mutants. Moreover, our confirmation that Mllt11 associates with α-tubulin in fetal brain lysates provides an explanation for the phenocopy of *Mllt11* mutants to those affecting tubulin isoforms. The dynamic expression of Mllt11 during UL CPN development also supports its role as a critical regulator of UL migration. Inefficient extension of neurites may also be attributed to impaired trafficking of organelles, ribosomes for local protein synthesis, or vesicles containing secreted cues and growth cone machinery to the axonal periphery (Gonzalez et al., 2016; Kennedy and Ehlers, 2006). Specifically, the potential Mllt11 interactor Myosin 10 we identified by proteomics is known to regulate radial migration of cortical neurons through its affect on the localization of N-cadherin (Lai et al., 2015), and has been implicated in tumor invasion through its role in process extension (Ropars et al., 2016). It is tempting to speculate that altered neuronal trafficking may be why *Mllt11* mutant UL neurons showed a progressive loss in the maintenance of an UL projection morphology and the transcription factor programs.

Loss of *Mllt11* from the superficial cortex altered levels of UL2/3 specific transcription factors Satb2 and CDP/Cux1, which function in establishment and maintenance of the corpus callosum as well as mature arborescent neuronal morphology and somal packing (Alcamo et al., 2008; Cubelos et al., 2015; He et al., 2021; Rodriguez-Tornos et al., 2016). Importantly, the loss of layer-specific morphology and identity has been demonstrated in postmortem brains from autistic humans (Casanova et al., 2013; Fujimoto et al., 2021; Stoner et al., 2014). These clinical data are reminiscent of the severe dysplastic appearance of superficial CPNs in the cortex of our conditional *Mllt11* mutants, revealed by the Golgi staining method. It is presently unclear whether the loss of UL-specific gene expression we observed in our *Mllt11* mutants, and tubulinopathies more generally, is primarily due to a disordered cytoskeleton, or through some other uncharacterized gene expression pathway functioning downstream of Mllt11. It is important to note that some neuronal subtypes exhibit altered expression of subpopulation-specific markers when cytoskeletal regulation is altered such that connections to target regions cannot be established (Hippenmeyer et al., 2005). This implies that there may be uncharacterized feedback mechanisms between establishment of connectivity and maintenance of expression of CPN subtype-specific transcription factors. Other potential roles for Mllt11 may include binding to HSPa8 and regulating protein export from the nucleus required for protein degradation (Li et al., 2014). HSPa8 was identified as a potential Mllt11 in our brain lysate proteomics, the meaning of which has yet to be explored. Yet another possible function for Mllt11 may include regulating the Wnt signaling via T cell factor 7, which in turn regulates CD44 to promote cell migration and metastasis (Li et al., 2018; Park et al., 2015). As ours is the first study to explore Mllt11 function in the developing brain, additional studies are needed to shed light on whether Mllt11 could be exerting its effect on transcriptional regulation directly, indirectly via its cytoskeletal-dependent interactions and intracellular trafficking, or both.

In light of this, we showed that the overexpression of *Mllt11* promoted migration into the cortical plate, consistent with a cytoskeletal regulatory function. This is also consistent with *in vitro* studies, which showed that the overexpression of *Mllt11* in cancer stem cell lines can promote their proliferation and invasiveness (Tse et al., 2017). Possible downstream effects of Mllt11-dependent cytoskeleton re-organization events may lead to favouring the symmetric divisions of organelles, nuclear components, and polarity protein complexes, all of which favour invasiveness of cells (Nance and Zallen, 2011; Piroli et al., 2019). On the other hand, the transcriptional regulation of Mllt11 by REST has been associated with promoting neuronal differentiation, with high REST expression correlating and low Mllt11 in undifferentiated neuronal tissue, and increasingMllt11 levels associated with decreased REST activity during terminal neuronal differentiation (Hu et al., 2015). This implies that there are potentially two discrete pathways through which Mllt11 regulates neuronal development: a cyctoskeletal regulatory mechanism, and a transcriptional regulatory mechanism.

Much of what is known about Mllt11 has been discerned from overexpression studies in immortalized cell lines and oncogenic clinical case studies, revealing pathogenic roles in cell process extension, increased invasiveness, and secretion of factors that promote efficient motility of cells (Chang et al., 2008; Li et al., 2006; Lin et al., 2004; Park et al., 2019; Tiberio et al., 2017; Tse et al., 2004). Oncogenesis and neurogenesis share commonalities at the level of cytoskeletal regulation of somatic and nuclear morphology, as well as extension of and trafficking along cytoplasmic processes, allowing for invasion of and migration through tissues. However, it is important to make the distinction between these pathological studies and our current findings on the role of Mllt11. Until the current study, the physiological role for Mllt11 was unclear. In all the published *in vitro* and oncogenic case studies, Mllt11 is aberrantly overexpressed in nonneuronal tissues either by itself or as a fusion with the chromatin remodelling protein Mll. Despite this, when we examined *Mllt11* expression during embryogenesis, we found it to be exclusively localized to developing neurons, and not in another other tissues, such as those deriving from mesoderm or endoderm (Yamada et al., 2014). The expression pattern of β-gal in the targeted *Mllt11* allele confirmed that it is normally exclusively expressed in developing neurons, with the highest levels reflecting UL cortical neurogenesis. Consistent with this, we now show that Mllt11 is required for proper migration, but not neurogenesis of UL CPNs. Furthermore, we also revealed that Mllt11 regulates neurite outgrowth and the maintenance of UL gene expression programs. The loss of *Mllt11* led to a reduction of interhemispheric connectivity *via* reduced crossing of fibres along the corpus callosum. We also characterized a severe dysplasia of CPNs in *Mllt11* cKO mutant neonatal brains; a phenotype found in severe neurodevelopmental disorders such as autism. Finally, we provide a possible mechanism of action for Mllt11 to link these phenotypes via its association with microtubules.

In summary, we investigated the role of Mllt11 in development of UL CPNs using a genetic knockout and labeling strategy to target the superficial cortex. Numbers and laminar distribution of UL CPNs were assessed over development, as well CPN neurite morphology. By combining *in vitro* and *in vivo* neurite outgrowth and morphology assays, we demonstrate that Mllt11 is required for neuronal invasion in the cortical plate, formation of mature dendritic morphologies characteristic of UL CPNs, and the extension of UL axons across the corpus callosum. Mllt11 interacts with the microtubule cytoskeleton and likely exerts its effect by altering cytoskeletal organization during development. Whether this occurs through the stabilization of cytoskeletal architecture or by trafficking of cellular machinery along neuritis will be the subject of future investigations.

## MATERIALS AND METHODS

### Animals

All animal experiments were done according to approved protocols from the IACUC at Dalhousie University. Mice (*Mus musculus*) carrying a null mutation in the Mllt11 gene were generated using embryonic stem (ES) cell clones obtained from the mouse knockout consortium project (UCDavis KOMP repository, *Mllt11tm1a(KOMP)Mbp*). The targeting construct is a “knock-out first, conditional second” approach, which inserts the gene encoding *β-galactosidase* (*β-gal*) exon 2, the protein-coding sequence of the *Mllt11* gene. Two independently targeted clones were injected into blastocysts, and the resulting chimeras were mated to BL/6 females to achieve germ-line transmission. Offspring were genotyped by PCR using the following primers: Wild Type F=5’-CGGTCCTGCCTTTGATTCTCAGC-3’ and R=5’-GCCTACTGCACAAGGTTCTTCTTGG-3’ (expected product size: 379bp), Mutant F=5’-GAGATGGCGCAACGCAATTAATG-3’ and R=5’-AAGCAGTATTTGCTTACTGGCCTGG-3’ (expected product size: 274bp). Heterozygotes (*Mllt11tm1a(KOMP)Mbp*/+) were maintained on a C57BL/6 background and crossed with *FlpO*^+/-^ (*B6.Cg-Tg(Pgk1-flpo)10Sykr/J*, 011065, The Jackson Laboratory) mice and converted to a conditional allele via germ line Flp recombinase expression. Resulting *Mllt11^Flox/Flox^* offspring were crossed with *Ai9 Rosa26TdTomato* (B6.Cg-*Gt(ROSA)26Sor^tm9(CAG-tdTomato)Hze^*/J, 007909, The Jackson Laboratory) reporter line to generate *Mllt11^Flox/Flox^; Rosa26TdTomato*^+/-^ or *Mllt11^Flox/Flox^; Rosa26TdTomato*^+/+^ offspring. *Cux2iresCre* mice used in this study to delete Mllt11 in developing UL neurons were previously described (Gil-Sanz et al., 2015; Yamada et al., 2015). *Cux2iresCre^+/-^* mice (*B6(Cg)*-*Cux2*<*tm1.1(cre)Mull*>/*Mmmh*, Mutant Mouse Resource and Research Center) were crossed with *Ai9* (*B6.Cg-Gt(ROSA)26Sor^tm9(CAG-tdTomato)Hze^*/J, 007909, The Jackson Laboratory) mice. The resulting *Cux2iresCre^+/-^;Rosa26TdTomato^+/-^* or *Cux2iresCre^+/-^;Rosa26TdTomato^+/+^* offspring were crossed with the *Mllt11^Flox/Flox^; Rosa26TdTomato^+/+^* line to conditionally knock out Mllt11. Offspring of these crosses were genotyped using the following primers: (Wild Type F=5’-CGGTCCTGCCTTTGATTCTCAGC-3’ and R=5’-GCCTACTGCACAAGGTTCTTCTTGG-3’ (expected product size: 379bp), Post Flp/Cre F= F=5’-CGGTCCTGCCTTTGATTCTCAGC-3’ and R=5’AAGCAGTATTTGCTTACTGGCCTGG-3’, and Cre F=5’GTTATAAGCAATCCCCAGAAATG-3’ and R=5’-GGCAGTAAAAACTATCCAGCAA-3’. We genotyped for the presence of TdTomato using primers F=5’ TACGGCATGGACGAGCTGTACAAGTAA-3’ and R=5’-CAGGCGAGCAGCCAAGGAAA-3 (expected product size: 500bp) and allelism was determined with primers F=5’-TCAATGGGCGGGGGTCGTT-3’, R=5’-TTCTGGGAGTTCTCTGCTGCC-3’, and R=5’CGAGGCGGATCACAAGCAATA-3’ (expected product size: 250pb for wild type and 300bp for mutant)

### GST protein expression, Mass Spectrometry, and Immunoprecipitation

#### GST-tagged protein expression

MLLT11 was cloned into pGEX-4T-2 vector for GST pulldown assays. BL21 competent cells (Sigma) were transformed with pGEX-4T-2 (GST) and pGEX-4T-2-MLLT11 (GST-MLLT11). Colonies were picked and incubated overnight in 2mL 2xYT media with ampicillin at 37°C with shaking. The following day, overnight culture was added to 150 mL 2xYT media with ampicillin and incubated at 37°C with shaking until OD_600_ reached 0.6. Protein expression was induced by adding IPTG (Invitrogen) to the culture to a final concentration of 0.1mM and shaken at 28°C for approx. 4 hours. Cells were pelleted by centrifugation and resuspended in 8mL NP-40 lysis buffer (50mM Tris pH 8.0, 150mM NaCl, 1% NP-40) with protease inhibitor cocktail (Sigma). Cells were nutated at 4°C for 10 mins, sonicated on ice (30s on/30s off, for 3 minutes total, level 6 intensity), and again nutated at 4°C for 10mins. Lysed cells were pelleted and supernatant collected (protein lysate).

#### Immobilization of GST or GST-MLLT11 bait protein

GST or GST-MLLT11 proteins were immobilized to Glutathione Sepharose 4B beads (Pierce). GST or GST-MLLT11 protein lysates were added to equilibrated 50% bead slurry and nutated at 4°C for approximately 3 hours. Beads with immobilized protein were collected, washed, and resuspended in PBS to make a 50% slurry.

#### GST pull-downs and Mass Spectrometry

Whole embryonic mouse brains were harvested at E15.5 and lysed in NP-40 lysis buffer with protease inhibitor cocktail (Sigma). Whole brain lysates from one embryo were added to either 6μg GST or 6μg GST-MLLT11 bead slurries and nutated overnight at 4°C. Beads with bound lysate were pelleted, washed, resuspended in sample buffer. Samples were heated for 15 mins at 37°C and run on an SDS-PAGE gel, then stained with Coomassie Blue (Pierce). Bands were cut from the SDS-PAGE gel and processed by Dalhousie’s Biological Mass Spectrometry Core Facility.

#### Co-immunoprecipitation

HEK293 cells were transfected with myc or myc-MLLT11 vectors using Lipofectamine™ 2000 (Invitrogen). 24 hours later, cells were lysed on ice with Tris-HCl lysis buffer containing 50mM Tris-HCl pH 7.4, 150mM NaCl, 1mM EDTA, 1% Triton X-100, and protease inhibitor cocktail (Sigma). Lysed cells were collected, centrifuged, and supernatant used for co-IP. Anti-c-Myc Agarose resin (Pierce) was used, as per manufacturer’s protocol, to co-immunoprecipitate myc or myc-MLLT11 and their binding partners. Proteins were eluted from the resin with 50mM NaOH, neutralized with 1M Tris pH 9.5, and added to non-reducing sample buffer for Western blot analysis.

### Western blots

E18.5 cortical protein samples were separated on 8% SDS-PAGE gels for 1hr at 120V and transferred overnight at 20V on to PVDF membranes (BioRad). Blots were probed with rabbit anti-tubulin β3 (1:1000, BioLegend), mouse anti-acetylated tubulin (1:20,000, Sigma), mouse anti-GAPDH (1:1000, Invitrogen), and rabbit anti-GAPDH (1:1000, Elabscience). Secondary antibodies were goat anti-rabbit HRP (1:5000, Invitrogen) and goat anti-mouse HRP (1:5000, Invitrogen). Blots were developed with Clarity Western ECL Substrate (BioRad) and imaged on a ChemiDoc™ Touch Gel Imaging System (BioRad). Band densitometry was done using Image Lab Software (BioRad).

### qPCR

To confirm the loss of Mllt11 RNA, E18.5 RNA was extracted from cortices of 3 genotypic conditional knockouts and 4 controls using the RNeasy Micro Kit (Qiagen). RNA was reverse transcribed to cDNA using the SuperScript™ II Reverse Transcriptase Kit (Invitrogen). qPCR reactions were carried out using the SensiFAST™ SYBR^®^ No-ROX Kit (Bioline) with the following primers for Mllt11 and internal control GAPDH: Mllt11 F=5’-GAACTGGATCTGTCGGAGCT-3 and R=5’-GCGCTCTCCAGAAGTTGAAG-3’, GAPDH F=5’-ACCACAGTCCATGCCATCAC-3’ and R=5’-TCCACCACCCTGTTGCTGTA-3’. The source for GAPDH primers was (Weng et al., 2014).

### Histology

Fetal brains were dissected out and fixed in 4% paraformaldehyde (PFA) for 4-8 hours, depending on embryonic stage, before being equilibrated in sucrose, embedded in Optimum Cutting Temperature (OCT) compound (Tissue-Tek, Torrance, CA), and cryosectioned at 12μm. Cortex morphology was assessed by DAPI staining. ß-Gal staining was performed on cryosectioned slides using the β-Gal Tissue Stain Kit (Millipore). Immunohistochemistry was conducted on E14.5-E18.5 brains as described previously (Iulianella et al., 2008). *Immunohistochemistry* was conducted using the following antibodies: rabbit anti-CDP/Cux1 (1:100; Santa Cruz), mouse anti-Satb2 (1:250; Abcam), rat anti-Bcl11b/Ctip2 (1:500; Abcam), rabbit anti-Tbr1 (1:200; Abcam), goat anti-Nestin (1:250; Santa Cruz), rabbit anti-Cleaved Caspase 3 (CC3, 1:500; Cell Signaling Technology), chicken anti-Tbr2 (1:500; Millipore), rabbit anti-Pax6 (1:500; Abcam), goat anti-Sox2 (1:200; Santa Cruz), mouse anti-NeuN (1:200; Millipore), rat anti-Tbr2 (1:200; eBioscience), mouse anti-Ki67 (1:25; BD Biosciences) and mouse anti-neurofilament 2H3 (1:200; DSHB, University of Iowa). Species-specific AlexaFluor 488-, 568-, 594-, and/or 647-conjugated IgG (1:1500; Invitrogen) secondary antibodies were used to detect primary antibodies.

*In situ hybridization* was performed on 30mm frozen sections obtained from E18.5, P7, P14, P21, and P28 Control and cKO (N=3/genotype) brains fixed overnight as previously described (Yamada et al., 2014) using an Mllt11 riboprobe (Chen et al., 2008; Toma et al., 2014). For *EdU birth dating studies*, dams were injected intraperitoneally with 30mg/kg body weight of EdU (Invitrogen) at E14.5 and E16.5 and E18.5 and sacrificed at E14.5 or E18.5. Sections were immunostained using the Click-It Kit according to the manufacturer’s protocol (Invitrogen). For *Golgi stains*, brains were harvested from mice at P28 and subjected to the FD Rapid GolgiStain Kit (FD Neurotechnologies, Inc.) as per manufacturer instructions. Crude sections were cut and mounted on slides with Permount Mounting Medium (Fisher Scientific). 4 brains were analyzed per genotype and 30 neurons per individual were analyzed.

#### Microscopy

Images were captured using a Zeiss AxioObserver fluorescence microscope equipped with an Apotome 2 structured illumination device, 10x, 20x, and a Hamamatsu Orca Flash v4.0 digital camera. ß-Gal and *Mllt11* in situ staining was captured using an upright Zeiss PrimoStar compound microscope with an ERc5s colour camera. Images were processed using Zen software (Zeiss, Germany) and Photoshop CS6 (Adobe, San Jose, CA).

### Primary cortical cell culture and immunocytochemistry

Cortices were microdissected from E18.5 embryos, digested in trypsin (Pierce), manually triturated and plated on 35mm well onto poly-D-lysine coated coverslips at a density of 150,000 cells for the neurite outgrowth assay or 500,000 for cellular localization experiments. Cells were plated in medium containing DMEM with 10% FBS and 1% penicillin/streptomycin and 4 hours after plating, media was completely removed and replaced with Neurobasal media containing B-27+ (Gibco/Invitrogen), 1% penicillin/streptomycin, and L-glutamine. Cells were cultured for 24 hours or 1 week in a 37°C incubator containing 5% CO2, then fixed for 10 minutes in 4% paraformaldehyde. Immunocytochemistry was conducted using the following antibodies: mouse anti-Tau (1:200; Abcam), rabbit anti-MAP2 (1:1000; Abcam), rabbit anti-Mllt11 (1:300, Abcam), rabbit anti-Tuj1 (1:1000, Biolegend), and mouse anti-acetylated α-tubulin (1:1000, Millipore Sigma).

### *In utero* electroporation and cDNA constructs

*In utero* electroporation was conducted using standard methodology under sterile surgical conditions (Saito, 2006). Endotoxin-free DNA was prepared according to the instructions of the manufacturer (Qiagen) and injected at 1.5 μg/μl into the telencephalic vesicles of embryos in time-staged pregnant females anesthetized under inhalable isoflurane (5L/min). A small incision was made on the ventral midline of anesthetized pregnant FVB females under a sterile field treatment. Single uteri containing the E13.5 fetuses were extruded and electric current was delivered across the fetal brains as 5 pulses for 50ms at 900ms intervals using tweezer-style electrodes linked to the pulse generator CUY21 Vivo SQ (Sonidel). The embryos were returned in the body cavity, the peritoneum was sutured and the skin was stapled. Experimental plasmids used were *Mllt11-ires-eGFP*, and control plasmids included *pIRES2-EGFP* (Clonetech) or pCIG (Addgene). To ensure comparable development staging, for each dam one uterus was electroporated with the experimental construct and the other with the control vector. Fetuses were allowed to survive for 2 days until E15.5 after which they were processed for cryosectioning to evaluate GFP expression and cortical layer development by immunostaining. N=3 Mllt11-eGFP and N=4 eGFP control fetal brains were analysed. Line graphs and statistical testing were conducted using Graphpad Prism V5.0d software, with results shown as mean ± SD. Statistical differences were determined with a Welch’s t-test.

### DiI tracing of callosal projections

Brains of E18.5 embryos were removed and embedded in 7% low gelling temperature agarose in DMEM medium with 1% penicillin/streptomycin. Embedded brains were crudely sectioned by hand at a thickness of approximately 1-2mm. Approximately 0.5 μL of DiI (Invitrogen) was injected into the cortical white matter tracts at rostral and caudal axial levels and incubated for 8 hours in Tyrode’s solution. Tissues were then fixed overnight at 4°C in 4% paraformaldehyde before incubating for 2 weeks at room temperature in PBS with 1% penicillin/streptomycin. Images were captured using Zeiss V16 Axiozoom stereomicrscope.

### Image sampling, quantification, statistics, and Sholl analysis

For analysis of immunostaining markers and EdU, counting frames (100μm x 100μm) were placed in a vertical strip along the somatosensory (S1) cortex with the first counting frame along the edge of the ventricle. At least 3 histological sections within the somatosensory cortex from 3-8 different animals were analysed for each immunostain, EdU or electroporated vector. Cells that were positively labelled for both DAPI and the marker were counted within each frame using ImageJ (FIJI) (Schindelin et al., 2012). Analysis of directionality and amount of radial glia stain in the E18.5 neocortex using Nestin stained images, captured with identical acquisition parameters, was assessed using features found in FIJI (Image J). The ‘directionality’ plugin was used to calculate the percentage of fibres in the image aligned in the same direction. Automatic thresholds were then applied and the area covered by Nestin staining was measured and expressed as a percentage of the total image area. From each of these measurements population averages of WT and mutant were compared. For analysis of primary cortical cell culture, neurites were traced and measured and Sholl analyses were conducted using the “simple neurite tracer” plugin (FIJI). At least 50 cells were counted per individual (n=3). Image montages were assembled using Photoshop CS6.

To ensure consistency among samples, cell counts were restricted to the presumptive S1 (somatosensory cortex) of the embryonic brain. In all experiments, 3-6 Mllt11 KO and control embryos were used for quantification analysis using the unbiased and systematic sampling method we described previously (Yamada et al., 2015). Counts for Ctip2, Tbr1, Cux1, Tbr2, NeuN, Sox2, Pax6, Ki67, CC3, and Satb2 are represented as line graphs, which represent the proportion of DAPI stained cells expressing those markers. For EdU, TdTomato, and DAPI distribution analyses, proportion of total stained cells expressing markers within each cortical bin was quantified to ensure counting methods were consistent with previous studies (Chen et al., 2008; Lawrenson et al., 2017; Reyes et al., 2019). For the electroporation experiments, 3-5 independent samples were analysed. Bar charts, line graphs, and statistical testing were conducted using Graphpad Prism V5.0d software, with results shown as mean ± SD. In all quantification studies, statistical differences were determined with Welch’s t-test (two tailed), or paired t-test in the case of Sholl analyses, with significance level set at P≤0.05 (*P≤0.05, **P≤0.01, ***P≤0.001, ****P≤0.0001).

## ACKNOWLEDGMENTS

A.I. lab gratefully acknowledges funding support for this project from the Canadian Institutes of Health Research (CIHR PJT-388914). We thank Sarah Whitehead, Emily Capaldo, Jessica Clark, and Di Shao for maintenance of mouse transgenic colonies, and Marina Gertsenstein at the Centre for Phenogenomics in Toronto, Canada, for assistance in generating chimeric mice. We thank Leanne Clattenburg for advice on GST pull down experiment.

## SUPPLEMENTARY FIGURE LEGENDS

**Figure S1: *Mllt11* targeting strategy and expression timeline.**

(A) Graphic representation of the targeting construct inserted into the *Mllt11* locus. *Mllt11* expression can be evaluated by ß-Gal staining due to the insertion of *lacZ* cDNA in the targeted allele. A conditional knockout allele can be garneted by the removal of the *lacZ* and selection cassette by germline Flp recombination. (B) A reference coronal section of a control brain with a boxed region to indicate the area sampled in panel (C) *Mllt11* expression in the cortex and hippocampus from P7 to P28. RNA levels declined in the cortex and were indistinguishable from background by P28. Expression in the hippocampus remained detectable throughout all stages evaluated. Scale bar: 100μm.

**Figure S2: *Mllt11* conditional knockout validation by qPCR and ISH.**

(A) Quantitation (q)PCR fluorescence levels of control and cKO cortices normalized to internal control GAPDH. (B) Fold change of *Mllt11* cDNA transcript levels was significantly decreased in cKO relative to control brains. (C-D) Images of ISH of *Mllt11* riboprobe on P7 control and cKO cortices (C) and hippocampi (D) showed decreased labeling in the superficial cortex, corresponding to the Cux2-expressing region. CP, cortical plate; RFU, relative fluorescence units. Welch’s t-test, (A-B) N=4 controls, 3 cKOs, (C-D) N=3. Data presented as mean ± SD. **P≤0.01. Scale bar: 100μm.

**Figure S3: *Cux2iresCre-driven* TdTomato expression in the pallial cortex and corpus callosum.**

(A-B) *Cux2iresCre* driven TdTomato reporter expression was highest in UL2/3 of the cortex (A) and the corpus callosum (B) at P7, corresponding to regions where the excision event occurs. Scale bar: 100μm.

**Figure S4: *Mllt11* is expressed in the developing cortical plate and *Mllt11* loss affects the laminar distribution of cells.**

(A) Coronal sections of the targeted *Mllt11* locus (which inserted a *lacZ* cDNA) showing *Mllt11* expression across four time points during cortical neurogenesis through β-Gal staining. At E14.5, β-Gal expression was most intense in the cortical plate (CP), corresponding to UL neurogenesis. By E16.5 β-Gal staining intensity shifted to the superficial cortex where UL CPNs were accumulating. (B-D) Body weight (B), brain weight (C), and brain weight as a percentage of body weight (D) showed no difference between control and conditional knockouts (cKOs) at E18.5. (E) cKO cortices were thinner than controls at E18.5. (F-H) Thinning of the cKO cortex was progressive, with thicknesses being comparable at E14.5 (F) but exhibited reduced thickness and reduced distribution of cells in the superficial cortex at E16.5 (G), which increased in severity by E18.5 (H). Total cell counts are shown as bar graphs. Line charts represent percentage of positive cells normalized to DAPI+ nuclei per 100 μm x 100μm bin. Welch’s t-test, (B-D) N=8, (F, H) N=5, (G) N=6. Data presented as mean ± SD. n.s, not significant. *P≤0.05; **P≤0.01. Scale bars: (A) 100μm (E) 100 μm (F-H) 50μm. Ctx, cortex; CP, cortical plate; LGE, lateral ganglionic eminence; MGE, medial ganglionic eminence; MZ, marginal zone; PP, preplate; SP, subplate; VZ, ventricular zone.

**Figure S5: Loss of *Mllt11* had no impact on programmed cell death.**

(A-B) Cortical slices at E14.5 (A) and E18.5 (B) showed no differences in the levels of cleaved caspase-3 (CC3) in cKOs relative to controls. (C) CC3 staining in the retrosplenial area at comparable levels, which normally has enhanced apoptosis, included as a control for CC3 staining. White arrowheads indicate positive labeling. N=3 controls, 5 cKOs. Scale bar: 50μm. VZ, ventricular zone.

**Figure S6: Neural progenitor populations were largely unaffected in *Mllt11* cKO mutants.** (A-C) Coronal cortical slices stained for Pax6 with IHC was unaltered in cKO cortices relative to controls at E14.5 (A) E16.5 (B) and E18.5. (D-F) Sox2 levels were largely similar between cKO and controls at E14.5 (D) E16.5 (E) and E18.5 (F). (G-I) Tbr2 expression was also largely unaltered in cKOs at E14.5 (G) and E16.5 (H), but showed a significant trend towards decreased levels normalized to DAPI+ nuclei immediately above the Tbr2+ progenitor domain at E18.5 (I). Line charts represent percentage of positive cells normalized to DAPI+ nuclei per 100μm x 100μm bin. Welch’s t-test, (A-C, F-I) N=4, (D-E) N=3. Data presented as mean ± SD. n.s, not significant. *P≤0.05 ; **P≤0.01; ***P≤0.001. Scale bar: 50μm for (A-I). VZ, ventricular zone.

**Figure S7: Progressive loss of UL CPN marker expression in *Mllt11* cKO mutants.**

(A-B) Coronal slices stained for Satb2 with IHC were similar between *Mllt11* cKO and controls at E14.5 (A) but displayed a tendency toward decreased numbers in upper bins and a downward shift at E16.5 (B). (C-D) CDP/Cux1 levels were largely normal at E14.5 (C) but began to decrease at E16.5 (D). (E-F) TdTomato levels were comparable between control and cKO at E14.5 (E) and E16.5 (F). (G-H) Tbr1 levels and localization were comparable between control and cKO at E14.5 (G) and E16.5 (H). (I-J) Ctip2 expression and localization were largely unaltered in cKO at E14.5 (I) and E16.5 (J). Line charts represent percentage of positive cells normalized to DAPI+ nuclei per 100μm x 100μm bin for (A-D) and (G-J) and percentage of total TdTomato+ cells per 100μm x 100μm bin for (E-F). Welch’s t-test, (A-D, G, I-J) N=4, (E) N=5 controls, N=6 cKOs, (H) N=4 controls, N=5 cKOs. Data presented as mean ± SD. n.s, not significant. *P≤0.05. Scale bar: 50μm for (A-J). VZ, ventricular zone.

**Figure S8: CPNs lacking *Mllt11* maintained neuronal identity.**

(A-B) Coronal cortical slices at E16.5 (A) and E18.5 (B) showing expression of NeuN was maintained though the cortical plate, but was compressed and shifted downward in cKOs relative to controls. (C) GFAP staining in the cortex showed comparable levels between cKOs and controls. Unaltered NeuN and GFAP levels suggested glial and neuronal fates are maintained in controls and cKOs. Welch’s t-test, N=3. Data presented as mean ± SD. *P≤0.05; **P≤0.01; ***P≤0.001, ****P≤0.0001. Scale bar: 50μm. VZ, ventricular zone.

**Figure S9: CPNs lacking *Mllt11* failed to fully invade cortical plate but do not acquire a DL identity.**

(A-B) Coronal cortical slices of control and cKO and close-up views (a’,b’) of mice injected with EdU at E14.5 (A-a’) or E16.5 (B-b’) and harvested at E18.5. Slices are co-labeled with DL CPN marker Ctip2 (A-B). Significantly less EdU+ cells invaded the cortical plate of cKO relative to control cortices. Welch’s t-test, N=3 (controls and cKOs). Data presented as mean ± SD. *P≤0.05. Scale bar: 50μm for (A-D). VZ, ventricular zone.

**Figure S10: Radial glial fibres were unaltered in *Mllt11* cKOs.**

(A) Coronal cortical slice at E14.5 showing expression of Nestin in controls vs. *Mllt11* cKOs. (B-D) Representative angle (B), area (C) and dispersion (D) showed no significant differences between controls and cKO. (D) Nestin expression in the cortex at E16.5. (E-G) Representative angle (E), area (F) and dispersion (G) showed no significant differences between control and cKO cortices at E16.5. (I) Cortical expression of Nestin at E18.5. (J-L) Representative angle (J), area (K) and dispersion (L) show no significant differences between control and cKO at E18.5, but dispersion (L) trended towards an increase in cKOs relative to controls. Welch’s t-test, (A-D) N=4, (E-H) N=5, (I-L) N=3. Data presented as mean ± SD. n.s, not significant. Scale bar: 50μm for (A, E, I). VZ, ventricular zone.

**Figure S11: Cajal-Retzius cells in Cortical Layer 1 are unaffected by *Mllt11* loss.**

(A-B) Pial sections at E14.5 (A) and E18.5 (B) showed comparable levels of expression of CR markers P73 and Reelin in Layer 1 of cKOs vs. controls. Total co-labeled cells are shown as bar graphs. Welch’s t-test, N=3. Data presented as mean ± SD. n.s, not significant. Scale bar: 50μm.

